# Linking live-cell imaging to single-cell sequencing using genetic barcoding

**DOI:** 10.64898/2026.10.07.757257

**Authors:** Micha Müller, Silke J.A. Lochs, Leonie Apperloo, Rupa Banerjee, Joana F. Marques, Carlota Davo-Martinez, Pim M. J. Rullens, Marta Moreno-Gonzalez, Vasilis Papageorgiou, Bernhard A. Kramer, Geert J.P.L. Kops, Jop Kind, Marvin E. Tanenbaum

## Abstract

Linking cellular phenotype to genotype and gene expression is a central goal in biomedical research. Live-cell imaging captures dynamic cellular phenotypes, whereas single-cell sequencing measures genome-wide molecular states, but pairing these readouts for the same cell remains challenging. Here we present Waldo, a genetically encoded single-cell barcoding strategy that can be read out by both live-cell imaging and single-cell sequencing to connect single cells across both modalities. Waldo barcodes are generated by combinatorial expression of synthetic cell-surface receptors, each detected by a cognate peptide probe coupled either to a fluorescent dye or a DNA barcode. Iterative fluorescent probe staining achieves large barcode diversity, while preserving fluorescence channels for phenotypic reporters. As a proof-of-principle, we use Waldo to link cell division phenotypes after chromosome missegregation to the karyotype and transcriptome of the same cells. Waldo provides a general strategy for connecting imaging phenotypes to genome-wide sequencing readouts in single cells.

## Introduction

How a cell’s phenotype—its dynamics, behavior and response to cues—relates to its genotype and gene expression is a central question in molecular cell biology. Live-cell imaging, from single molecules to whole cells and tissues, has transformed our understanding of cell behavior, while a wide range of single-cell sequencing approaches now allow gene regulation to be studied in depth in single cells^1^. Each of these modalities is transformative in its own right, but applying both to the same single cell and linking the measurements has remained a major challenge. Matching a phenotypic readout to a gene expression readout in the same cell could provide a mechanistic understanding of how gene expression shapes cell behavior and can reveal the genetic basis of single-cell functional heterogeneity^2^. Combining these approaches lies at the heart of the emerging field of single-cell phenomics, which seeks to pair omics measurements with functional, phenotypic and imaging-based readouts in individual cells^3^.

A number of approaches already connect imaging and sequencing methods, based on a variety of different principles: physical capture of imaged cells^4^ or subcellular biopsies^5^, microfluidic isolation^6,7^, photolabeling of cells and isolation by fluorescence activated cell sorting (FACS)^8,9^, photoselective sequencing^10^, *in situ* imaging-based transcriptomics^11,12^ and spatial barcoding followed by sequencing^13,14,15,16^. While each of these approaches is powerful and has specific applications, none of these approaches allow linking of high-resolution, live-cell imaging of large numbers of cells *in situ* to single-cell sequencing-based readouts, leaving a critical gap in scalable approaches for relating genotype to phenotype in individual cells.

Here, we present Waldo (named after “Where’s Waldo?”, in which an individual must be found in a crowd by their distinctive clothing^17^), a genetically encoded barcoding method that can be read out in individual cells during both live-cell imaging and single-cell sequencing. The fluorescent barcode is read out iteratively, allowing, in theory, infinite barcode complexity. To establish a proof-of-concept, we used Waldo to study the early effects of chromosome missegregation during cell division. By linking live-cell imaging, single-cell RNA-sequencing and single-cell karyotype sequencing using Waldo, we uncover how cell division errors and subsequent micronucleus formation are linked to daughter cell karyotype and transcriptome of individual cells. More broadly, Waldo provides a versatile and widely applicable tool for linking live-cell imaging to single-cell sequencing, opening new avenues for studying a wide range of biological questions across diverse fields.

## Results

### Combinatorial expression of genetically encoded synthetic receptors to create single-cell barcodes

*The Waldo method in brief:* To combine live-cell imaging with single-cell sequencing, single cells are barcoded by combinatorial expression of a random subset of synthetic receptors. Cells are first analyzed by live-cell imaging to assess single-cell phenotypes, after which receptor expression is read out iteratively using fluorescently labeled probes, each binding to a unique synthetic receptor (Fig. 1a). After barcode readout, the cells that were analyzed by live-cell imaging are photoactivated, stained with DNA-labeled probes and recovered by FACS. Single-cell sequencing libraries are then prepared and the RNA and/or DNA is analyzed by single-cell sequencing. Finally, receptor expression of each sequenced cell is determined through quantitative analysis of DNA probe levels or through receptor mRNA sequencing, and barcodes are compared between live-cell imaging and single-cell sequencing to link cellular phenotype to gene expression.

**Fig. 1.**
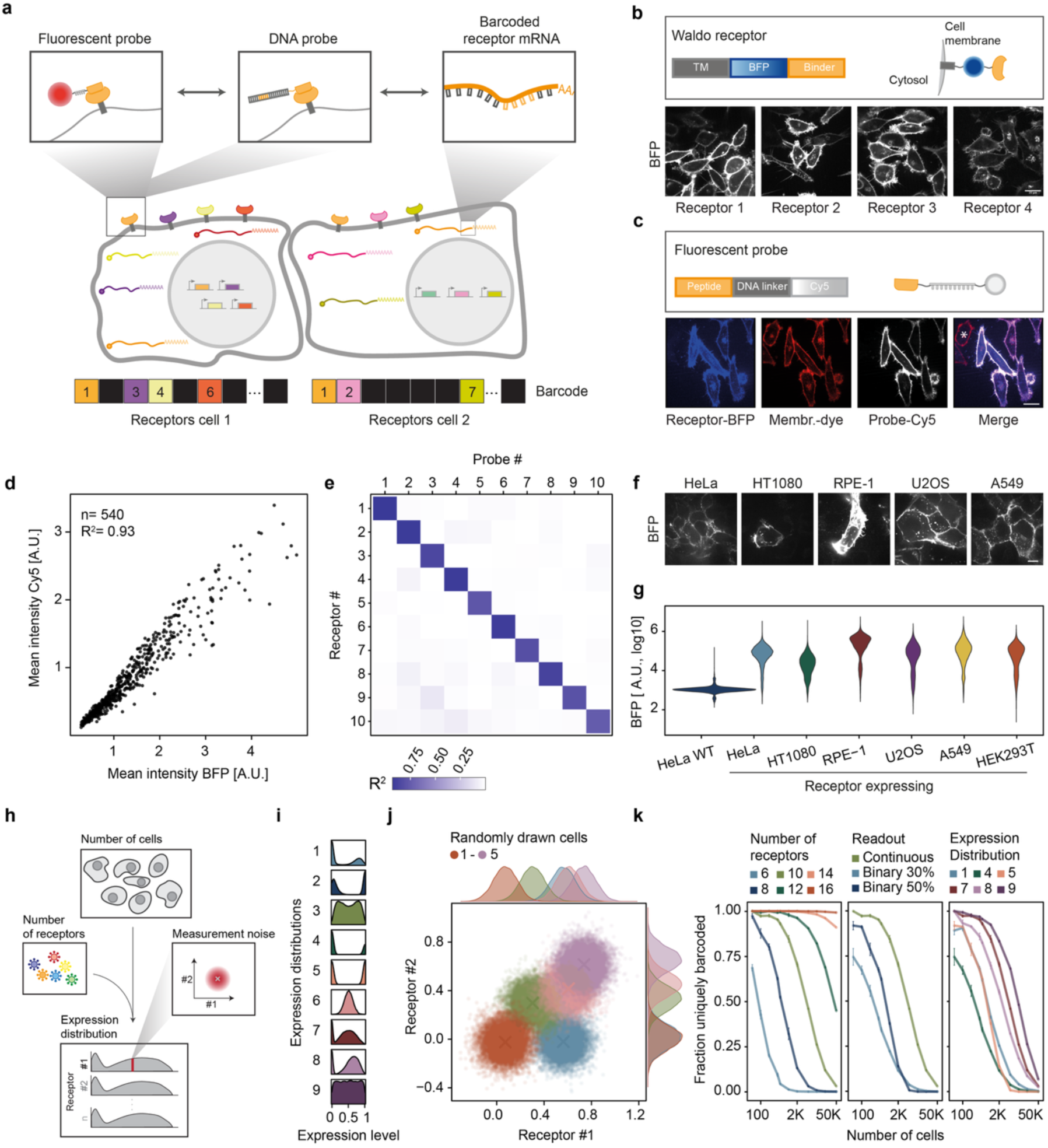
Combinatorial expression of genetically encoded receptors to create single-cell barcodes. **a**, The Waldo barcoding strategy. Barcodes are formed by combinatorial expression of cell-surface receptors and read out either by imaging, using fluorescent probes that bind the receptors, or by sequencing, using either DNA probes or barcodes in the receptor-encoding mRNAs. **b**, Top, receptor construct: transmembrane (TM) domain, blue fluorescent protein (BFP) and a probe-binding domain. Bottom, four Waldo receptors expressed in HeLa cells, each showing correct localization to the cell surface. **c**, Top, fluorescent probe: a peptide, a single-stranded DNA linker and a Cy5 dye. Bottom, receptor-expressing cells (receptor-BFP, blue) labelled by the probe (Cy5, white); non-expressing neighbors are labelled by membrane dye (red) but lack BFP and probe signal (indicated with asterisk). **d**, Single-cell receptor-BFP versus probe (Cy5) intensity for receptor–probe pair 1; n and R^2^ shown. **e**, Orthogonality matrix of all receptor–probe combinations; color encodes the R^2^ of BFP versus Cy5 intensity per pair (scale at bottom), with diagonal signal indicating cognate-only binding. Cells per comparison ranged from 244 to 848 (median 472). **f**, Representative images showing correct membrane localization of a Waldo receptor (BFP) in HeLa, HT1080, RPE-1, U2OS and A549 cells. **g**, Receptor-BFP fluorescence across indicated cell lines measured by flow cytometry; wild-type Hela cells without receptor expression (HeLa WT) were used as a control. **h**, Parameters varied in the barcode-diversity simulations: number of cells, number of receptors, receptor expression distribution and measurement noise. **i**, The nine simulated expression distributions: bimodal (1, 2, 3, 7, 8), binary (4, 5), Gaussian (6) and uniform (9). **j**, Simulated expression levels for two receptors for five cells drawn from distribution 8 (panel **i**). Crosses mark true expression; surrounding points and marginal densities show simulated measurement noise. Cells are colored from red (cell 1) to purple (cell 5). **k**, Fraction of uniquely barcoded cells. Left, effect of receptor number (distribution 8, noise 0.1 s.d.). Middle, effect of readout type (continuous, or binary with 30% or 50% positive cells; 10 receptors, noise 0.1 s.d.). Right, effect of expression distribution (10 receptors, noise 0.1 s.d.). Lines and points show the mean and error bars the s.e. of three simulation repeats. Scale bars, 20 µm (b, c, f).

To create the genetically encoded barcode that can be read out by both imaging and sequencing, we designed a set of synthetic cell-surface receptors that can be labeled by fluorescent or DNA-barcoded peptide probes and can be expressed in single cells in different combinations (Fig. 1a, Suppl. Table 1). Each receptor consists of a transmembrane domain, a blue fluorescent protein (BFP) and a unique extracellular domain that specifically binds a cognate peptide. These synthetic receptors localized robustly to the cell surface when expressed in human cells (Fig. 1b). For the imaging barcode readout, each receptor is detected by the cognate fluorescent peptide probe (fluorescent probe), consisting of a short (25 amino acids) peptide that binds the receptor’s peptide-binding domain. The peptide is linked to a Cy5 dye through a short (30 nt) single-stranded DNA linker, which enables rapid dye removal by a DNase after barcode receptor readout, see below (Fig. 1c). Successful conjugation of the DNA-dye fusions to peptides could be verified by gel electrophoresis (Suppl. Fig. 1a). When added to cells, the fluorescent probe specifically labeled receptor-expressing cells, whereas neighboring non-expressing cells show no probe binding (Fig. 1c). Importantly, the probe signal intensity scaled linearly with receptor expression (as assessed by BFP fluorescence) across single cells (Fig. 1d, R^2^ = 0.93). This quantitative relationship confirms that fluorescent probe labeling intensity accurately reports on receptor expression levels. Therefore, *expression level* of each receptor, not merely its presence or absence, can be used to encode information to increase the diversity of single-cell barcodes.

We next performed a number of control experiments to validate different parts of the Waldo approach. First, comparing the full set of receptor-probe combinations showed that probe-receptor binding was specific to cognate pairs, with negligible cross-reactivity between non-cognate pairs (Fig. 1e). Second, a Waldo receptor construct transduced into a panel of human cell lines, including HeLa, HT1080, RPE-1, U2OS, A549 and HEK293T, consistently localized to the cell surface, with comparable expression levels across all lines (Fig. 1f,g), indicating that Waldo is not restricted to a particular cell type. Third, in cells transduced with three receptors carrying different fluorescent proteins, expression levels of the three receptors were uncorrelated among different cells (Suppl. Fig. 1b,c), confirming that receptors are delivered to cells independently. Finally, plotting probe intensity against receptor-BFP intensity across the full dynamic range showed that measurement noise was independent of expression level (Suppl. Fig. 1d,e) and approximately Gaussian (Suppl. Fig. 1f), so a receptor’s expression level can be estimated with the same confidence whether expression is low or high.

We next asked how many cells could be uniquely barcoded by Waldo, and which parameters set maximum barcode diversity. We simulated receptor expression levels and quantified barcode diversity by determining the fraction of cells with a uniquely distinguishable barcode within the simulated cell population (see Methods; Fig. 1h,i,j). Varying the number of cells and receptors, as well as the expression distribution and measurement noise, we found that barcode diversity increased with the total number of receptors introduced in the population, as expected, and was strongly enhanced by inclusion of receptor expression levels as a barcode parameter compared to a binary (present/absent) readout. Broad expression distributions further increased diversity (Fig. 1k). With 10 receptors, a readout noise of 0.1 s.d. and a broad bimodal expression distribution (e.g. distribution 8 in Fig. 1i, most similar to our experimental expression distribution, see Fig. 1g), barcodes were sufficiently diverse to uniquely identify approximately 200 cells. The number of uniquely barcoded cells could be increased to >10,000 by inclusion of 4 additional receptors. These simulations show that combinatorial receptor expression can barcode large numbers of cells, and that barcode diversity increases sharply with the number of receptors, providing a route to scale barcode diversity further.

### Imaging of Waldo barcodes through iterative receptor staining of live cells

Previous efforts to fluorescently barcode single cells have relied on combinations of spectrally distinct fluorescent proteins^18,19^. While such multi-color barcoding approaches allow rapid barcode readout, this approach is limited in its maximal barcode diversity, as the number of barcode elements (e.g. fluorescent proteins) that can be discriminated is limited by the spectral overlap of different fluorescent proteins. To overcome this limitation, iterative barcode readouts have been explored in recent years^20,21^. However, these methods have relied on cell fixation and harsh chemical treatment, both of which degrade the quality of downstream sequencing libraries. We therefore developed an iterative imaging readout compatible with live cells (Fig. 2a, Methods); in each round, cells are stained with one or more fluorescent probes and imaged to read out receptor expression in each cell, after which the fluorescence associated with the probe is removed by a single-stranded-DNA-specific nuclease (ssDNase) that cleaves the DNA linker connecting the peptide to the Cy5 dye.

**Fig. 2.**
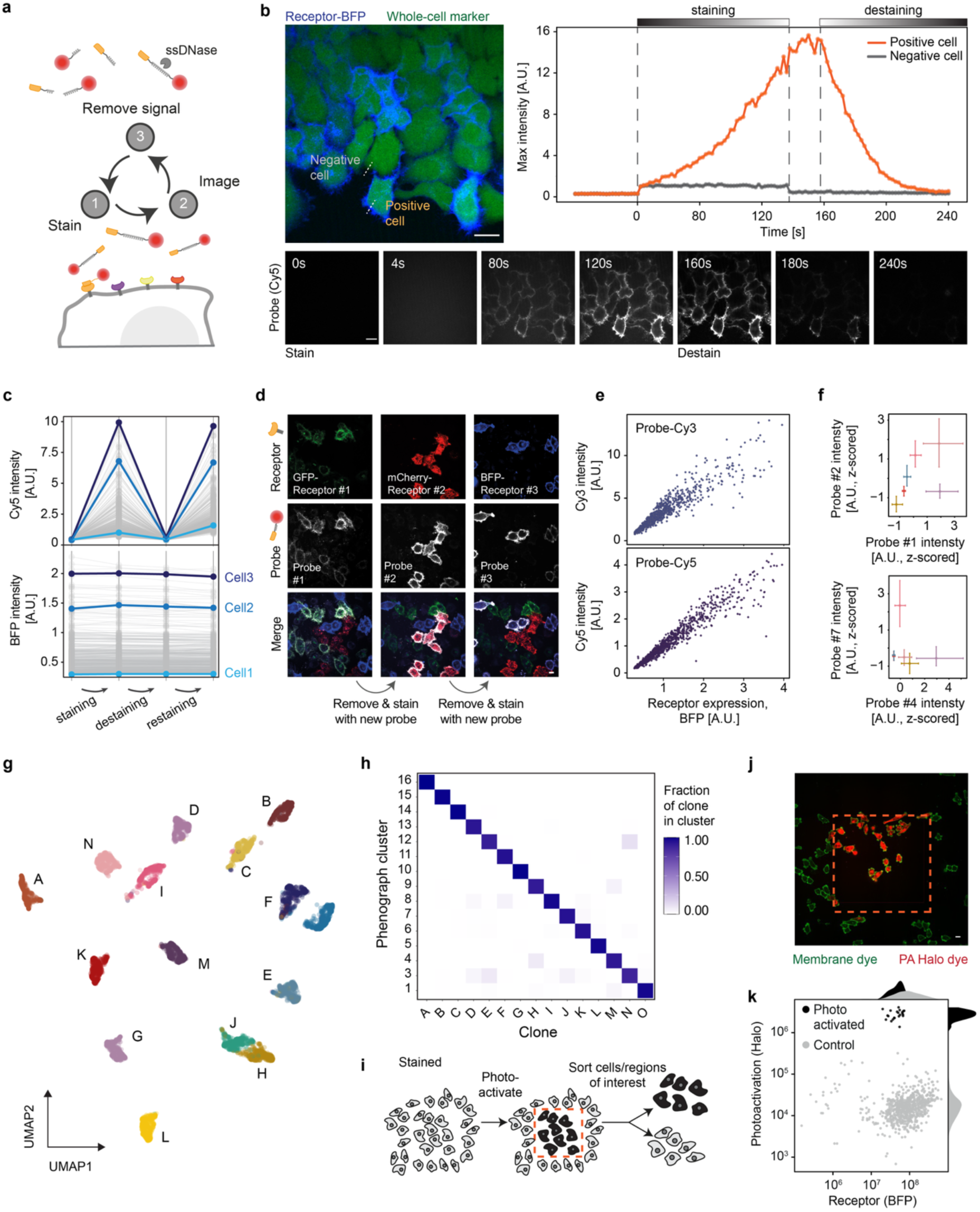
Imaging of Waldo barcodes through iterative receptor staining of live cells. **a**, Iterative readout of Waldo receptors: each round cells are stained with fluorescent probes, imaged, after which signal was removed using a single-stranded-DNA-specific nuclease (ssDNase) that cleaves the ssDNA linker that connects the peptide to the dye. **b**, Probe staining and destaining of live cells over time. Left, receptor-expressing cells (receptor-BFP, blue) and a whole-cell marker (green), with a receptor-positive and a receptor-negative cell indicated. Right, maximum probe (Cy5) intensity over time for the receptor positive (orange) and negative (grey) cell. Bottom, probe (Cy5) images of the same field at the indicated times (0–240 s). **c**, Probe (Cy5, top) and receptor (BFP, bottom) intensity of single cells over staining, destaining and restaining rounds; three cells with different receptor expression are highlighted (Cell1–Cell3). Receptor-BFP is stable, whereas probe signal cycles. **d**, Sequential multiplexed readout of three receptors in different cells (GFP-Receptor #1, green; mCherry-Receptor #2, red; BFP-Receptor #3, blue). Each round cells were stained with one cognate probe (receptor, top; probe, middle; merge, bottom) before removal and restaining. Each probe labels only its cognate receptor. **e**, Single-cell receptor (BFP) versus probe intensity for probes coupled to Cy3 (top, n=967 cells) or Cy5 (bottom, n=1,020 cells). **f**, Barcode intensities for selected probe pairs among individual cells belonging to different monoclonal cells lines expressing different receptors. Each cross shows the mean and 5th–95th percentile in probe intensity. **g**, UMAP of single-cell barcode intensities, colored by clone of origin (15 clones, A–O; subsampled to 440 cells per clone). **h**, Concordance between unsupervised clustering and clone of origin; color shows the fraction of a clone’s cells (A–O) per Phenograph cluster (1–16), with the diagonal indicating agreement. **i**, Schematic of selective photoactivation approach: cells stained with the photoactivatable HaloTag dye PA-JF646 are photoactivated within a selected region for subsequent isolation by FACS. **j**, Imaging after photoactivation (membrane dye, green; photoactivatable HaloTag dye PA-JF646, red); the dashed box marks the photoactivated region. **k**, FACS analysis of receptor (BFP) expression versus photoactivated PA-JF646. Photoactivated cells (black) clearly separated from non-activated cells. Scale bars, 20 µm (b, d, j).

Removing signal between rounds allows the same fluorescence channel to be reused, such that the number of receptors that can be read out is no longer limited by the number of available fluorescence channels. An additional advantage of reading out the barcode in a single fluorescence channel is that the remaining fluorescence channels are free for imaging complex cellular properties through multi-color live-cell imaging.

To assess the quality of the iterative barcode readout approach, we characterized the staining and destaining kinetics and efficiency of live cells (Fig. 2b, Suppl. Video 1). Within approximately 2-3 minutes of fluorescent probe addition, probe labeling of receptor-expressing cells was saturated (Fig. 2b). Signal removal by the ssDNase was even faster, returning to baseline fluorescence within ∼1.5 minutes. This labeling speed is facilitated by the small size of the probe and the cell-surface exposure of the receptor. Repeating iterative staining and destaining over two rounds at sub-saturating probe concentrations (to allow restaining of the same receptor) revealed that the probe signal recovered to the same intensity upon restaining (Fig. 2c), demonstrating that the nuclease is washed out efficiently to allow for the next round of staining. As an additional control for the specificity of the iterative barcode readout, we generated three cell lines, each expressing a different Waldo receptor fused to a different fluorescent protein (GFP, mCherry or BFP). Cell lines were then mixed, and barcode readout was performed for all three receptors using the iterative staining procedure. All three receptors were robustly and specifically labeled by their cognate probe (Fig. 2d), confirming that the iterative readout functions well.

While reading out the barcode in a single fluorescence channel allows live-cell multi-color phenotyping, barcode readout can also be performed in multiple fluorescence channels simultaneously using probes with spectrally distinct dyes. Such multiplexed readout reduces the number of barcode readout rounds and thus accelerates barcode readout. As a proof of concept, we generated both Cy3- and Cy5-coupled probes and confirmed that the quantitative readout was possible using both dyes (Fig. 2e, Suppl. Fig. 2a). This multiplexing capacity could be extended further with additional dyes or spectral-unmixing approaches^22^ to further enhance barcode readout speed.

To assess the precision and reproducibility in barcode readout, we generated 15 different monoclonal cell lines, each expressing a subset of Waldo receptors (Suppl. Fig. 2b, Methods). Because all cells within a clone carry the same genetic barcode, comparing cells within and between clones tests how accurately the fluorescent barcode can be read out. Probe intensity readout for many cells from each clone revealed that cells within a single clone showed similar probe staining intensities that could be readily distinguished from probe intensities of cells from different clones (Fig. 2f). This was further confirmed by reducing the full barcode space to two dimensions using UMAP, where cells from different clones separate into well-defined groups based on probe staining intensities (Fig. 2g). Unsupervised clustering of single-cell barcodes identified one cluster (cluster 2, containing ∼5% of the analyzed cells) corresponding to dead cells stained non-specifically by all probes; these were excluded from further analysis (Suppl. Fig. 2c). The remaining clusters predominantly represented cells of the same clone of origin (Fig. 2h). Together, these results show that cells can be confidently assigned to their clone of origin from the fluorescent barcode alone, confirming that the iterative imaging readout is robust and specific.

In a live-cell imaging experiment, the imaged cells typically represent only a small subset of all cells in the dish, yet all cells carry a Waldo barcode. Sequencing non-imaged cells would waste sequencing reads on cells that cannot be matched to any imaging data and would increase the pool of cells that individual sequenced cells need to be matched with, increasing the likelihood of incorrect matches. To allow selective isolation of the cells that had undergone analysis by imaging, we incorporated a photoactivation step at the end of live-cell imaging and barcode readout to mark all the cells in the field of view during live-cell imaging for later isolation by FACS (Fig. 2i). Photoactivation was performed in the same fluorescence channel as barcode readout so as not to interfere with live-cell phenotyping. To achieve this, cells are stained after barcode readout with a photoactivatable far-red HaloTag dye (PA-JF646,^23^), which we found, labels cells non-specifically at high concentrations (0.5 µM), eliminating the need to create cell lines that express a photoactivatable fluorescent protein or HaloTag. After staining, cells within a selected region were photoactivated and isolated by FACS, with photoactivated cells showing a strong increase in fluorescence compared to non-activated cells (Fig. 2j,k). Together, these results demonstrate that the iterative imaging readout provides fast, specific and reproducible barcode detection in live cells, and that photoactivation followed by FACS enables selective isolation of the imaged cells, providing an optimal starting point for downstream single-cell sequencing.

### Readout of Waldo barcodes in single-cell sequencing

To read out Waldo barcodes in single-cell sequencing, we developed two complementary approaches: quantification of barcoded DNA-conjugated probes (’DNA probes’) bound to Waldo receptors, and direct readout of Waldo receptor mRNA expression (Fig. 3a, Suppl. Fig. 3a, Suppl. Table 2).

**Fig. 3.**
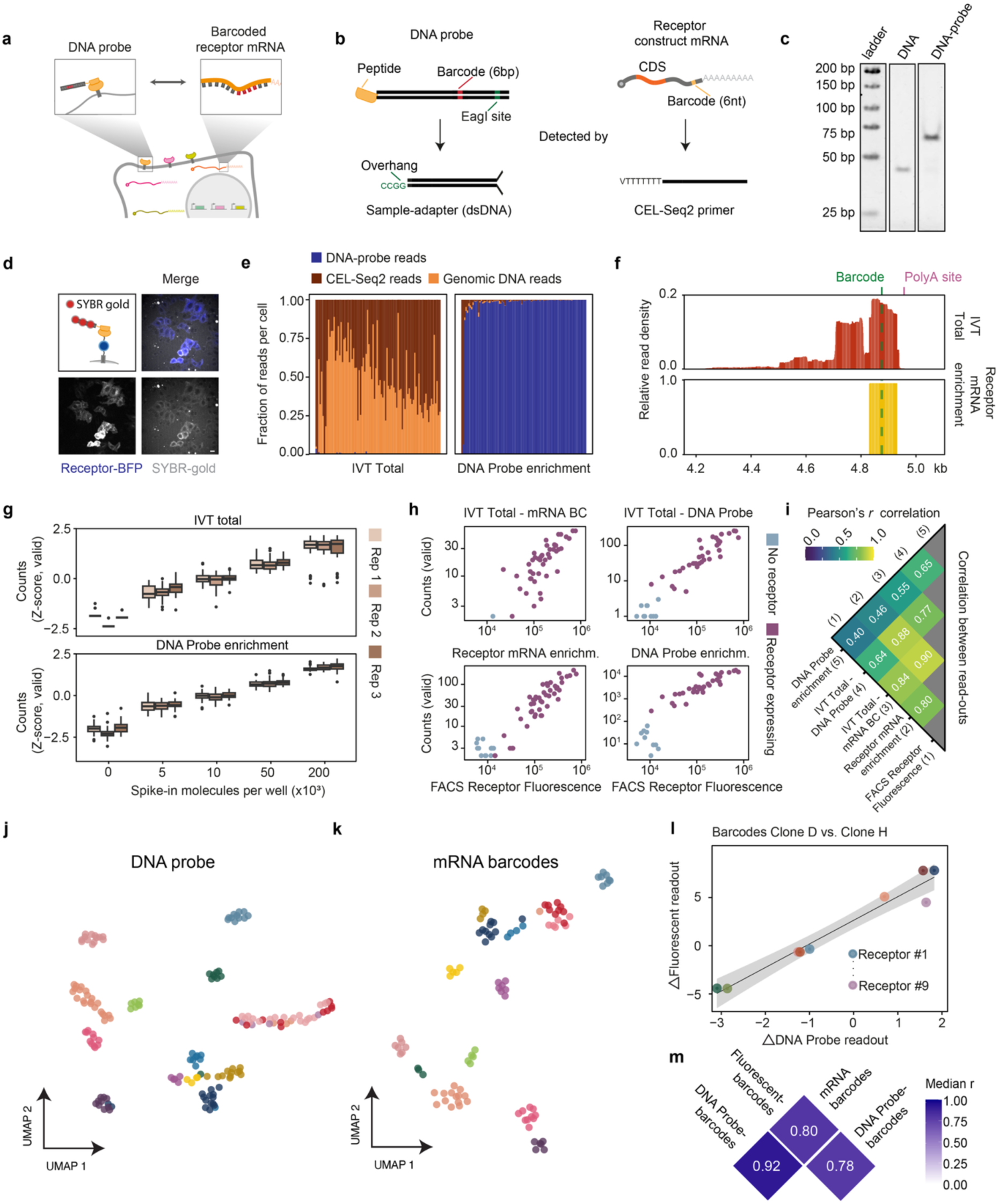
Sequencing of Waldo barcodes in single cells. **a**, The two single-cell sequencing readouts of Waldo barcodes: via DNA probes bound to the receptors, or via a barcode introduced in the receptor-encoding mRNAs. **b**, Detected molecules and their library adapters. The DNA probe (left) carries a peptide (orange) conjugated to a dsDNA oligo with a probe-specific 6-bp barcode (red) and an EagI site (green) for sample-adapter ligation. The receptor mRNA (right) carries the coding sequence (CDS, dark orange) and a 6-nt barcode (light orange) in the 3′ UTR near the poly(A) tail, captured by CEL-Seq2 poly(T) primers. **c**, In-gel fluorescence of the uncoupled dsDNA oligo (DNA, left) and the dsDNA oligo conjugated to the peptide (DNA probe, right). **d**, DNA probe staining of receptor-expressing cells (blue), visualized with SYBR Gold (grey). Merge is shown (top right). **e**, Fraction of read types per cell (each column, one cell) for the IVT total library (left) and the DNA probe-enriched library (right): DNA probe reads (blue), CEL-Seq2 reads (dark red), genomic-DNA reads (orange). **f**, Aligned-read density across the 3′ end of the receptor mRNA for the total IVT (top) and receptor-mRNA-enriched (bottom) libraries; green dashed line, barcode position; pink dashed line, poly(A) site. **g**, z-score-normalized valid counts for variable spike-in amounts (number of spike-in molecules per well) across three replicates (Rep 1–3); top, IVT total; bottom, DNA probe enrichment. Boxes show median and IQR, whiskers 1.5× IQR, points outliers. n=187–217 cells per replicate (median 203). **h**, Valid counts versus FACS receptor fluorescence for the mRNA barcode or DNA probe (coming from IVT Total or barcode enrichment libraries), a representative replicate is shown; receptor-expressing cells, purple; no-receptor controls, light blue. n=46 (DNA probe, IVT total), n=46 (DNA probe, enrichment library), n=44 (mRNA barcode, IVT total) and n=60 (mRNA barcode, enrichment library) cells. **i**, Pearson’s *r* between read counts (mRNA barcode or DNA probe, either from IVT or enrichment libraries) and FACS receptor fluorescence. Values represent the mean across three replicates (scale at top). **j**, UMAP of DNA probe barcode space, colored by clone of origin (n=164 cells). **k**, UMAP of mRNA barcode space, colored by clone of origin (n=113 cells). **l**, Relative expression of different receptors is similar in imaging (fluorescent readout) and sequencing (DNA probe readout) readouts. Relative receptor expression for cells from clones D and H is shown. Each point represents relative expression of one of nine receptors (Receptor #1–#9), showing the clone D-versus-H difference in median standardized signal; black line, linear fit; grey band, 95% confidence interval. **m**, Agreement between the DNA probe, fluorescent and mRNA barcode readouts across all clone pairs. For each clone pair, per-receptor differences in median standardized signal were correlated (Pearson’s *r*) between readouts; tiles show the median *r* across clone pairs. Scale bar, 20 µm (d).

#### DNA probe barcode readout approach

To read out Waldo barcodes using the DNA probes, Waldo peptides are conjugated to a double-stranded DNA molecule, instead of the fluorescent dye used for the imaging readout. Each DNA molecule carries a probe-specific 6-bp barcode, unique molecular identifiers (UMIs), PCR primer sites, and an EagI restriction site that allows ligation of a cell-specific sample adapter, similar to the approach used in MAbID^24^ (Fig. 3b; Methods). DNA–peptide coupling was achieved using the same conjugation chemistry as for fluorescent probes and was confirmed by gel-shift analysis (Fig. 3c; Methods). To verify that DNA conjugation did not impair probe binding to Waldo receptor, we stained DNA probes with SYBR gold, a dye that labels nucleic acids, and incubated DNA probes with receptor-expressing cells, which resulted in clear fluorescent labeling of receptor-positive cells (Fig. 3d).

We also confirmed that prior fluorescent probe labeling of cells did not prevent subsequent DNA probe binding to the same cells (Suppl. Fig. 3b). Re-labeling of the same receptors is likely possible because of turnover of fluorescent probe-receptor interactions over time.

Before staining cells with DNA probes, cells also need to be detached from the imaging plate, but we found that standard trypsin-based cell detachment cleaves the extracellular domain of Waldo receptors, abolishing probe binding to Waldo receptors (Suppl. Fig. 3c). Cell detachment using Accutase or enzyme-free dissociation, however, retained functional probe binding (Suppl. Fig. 3c; Methods). After cell detachment and DNA probe staining, single cells were sorted by FACS and processed using an established plate-based library preparation protocol compatible with CEL-Seq2, scDam&T-seq, and MAbID^24,25,26^ (Suppl. Fig. 3a; Methods). During processing, EagI digestion and adapter ligation—required to append cell-specific barcodes to the DNA probes—also generates adapter-ligated genomic DNA fragments, which are co-captured alongside cellular mRNA (Suppl. Fig. 3a). DNA probe barcodes were then selectively amplified by PCR, yielding a library composed almost exclusively of DNA probe reads, hereafter referred to as the “DNA Probe enrichment” library (Fig. 3e).

#### Receptor mRNA barcode readout approach

Waldo barcodes can also be read out through introduction of a unique 6-nt barcode inserted into the 3ʹ UTR of each receptor construct (Fig. 3b). CEL-Seq2^25^ is a polyT-based scRNA-seq method that generates 3ʹ-biased reads. However, library preparation uses random hexamer priming, so reads are distributed across the 3ʹ end of transcripts and do not always cover the barcode insertion site. To ensure consistent coverage of the receptor-specific barcode in the mRNA reads, we designed targeted reverse transcription (RT) primers that hybridize upstream of the barcode and cover the barcode insertion site. This targeted enrichment step increased the fraction of receptor-mRNA reads within the library (Fig. 3f), improving barcode detection sensitivity. This approach is hereafter referred to as the “Receptor mRNA enrichment” library.

Because both DNA probes and receptor mRNAs can be captured in the same single-cell library preparation, the two barcode readout approaches can be integrated in a single barcode readout. Overall, the Waldo approach yields three parallel single-cell libraries: IVT amplification of all captured material (“IVT total”, containing CEL-Seq2 mRNA, genomic DNA, and DNA probe reads), targeted receptor-mRNA enrichment (“Receptor mRNA enrichment”), and direct PCR of DNA probe (“DNA Probe enrichment”) (Suppl. Fig. 3a). Combining different barcode readout strategies may increase the precision of barcode readout, as each library contributes independent barcode observations. The readouts can, however, also be used independently: when barcode information is needed to match cells but mRNA sequencing is not available, the DNA probe PCR library can be used for barcode readout; conversely, when only mRNA sequencing is available, the receptor-mRNA readout can still provide barcode information for cell matching.

For Waldo to reliably link imaging and sequencing data, the sequencing-based barcode readout must quantitatively reflect receptor expression levels in single cells. As a first test of the quantitative nature of barcode readout in sequencing, barcoded spike-in DNA molecules were added at defined concentrations. Spike-ins were recovered in proportion to their input, confirming quantitative probe detection in the DNA probe libraries (Fig. 3g). To further assess how accurately the sequencing-based readouts reflect receptor expression levels in single cells, we compared barcode read counts for the different modes of barcode readout with receptor-BFP fluorescence for the same cells, as determined by FACS. Both DNA probe and receptor-mRNA barcode read counts showed a good correlation with BFP fluorescence (Fig. 3h), confirming the quantitative nature of the sequencing-based barcode readout. Receptor-mRNA barcodes showed the strongest correlation with FACS-based receptor BFP fluorescence (Pearson r=0.8-0.84). Overall correlations between BFP levels and PCR-based DNA probe reads were also high for receptor-expressing cells (Fig. 3h), although receptor-negative cells deviated from this linear correlation for this sample, somewhat reducing the Pearson correlation values (Fig. 3h,i). Together, these results demonstrate that both approaches to measure barcodes by sequencing quantitatively report on receptor levels in single cells.

Finally, to assess the reproducibility in sequencing barcode readout, we turned to the clonal cell lines, each expressing a subset of Waldo receptors which we already analyzed by imaging-based barcode readout (see Fig. 2f-h). As with the imaging-based readouts, cells from the same clone clustered strongly together in UMAP space when barcodes were read out by sequencing, both for the mRNA and DNA probe measurements (Fig. 3j,k). To determine whether imaging and sequencing yield concordant receptor expression levels, we compared the relative expression of each Waldo receptor between the two modalities across the different clones. Pairwise comparison of all clones showed that relative expression of different receptors was highly consistent between the imaging- and sequencing-based readouts (Fig. 3l,m), confirming that the Waldo method allows barcode readout and matching across both modalities.

### Studying immediate responses to chromosome missegregation using Waldo

As a proof-of-concept that Waldo can match imaging-based cellular phenotypes to genotype and gene expression, we applied Waldo to study the immediate cellular response to chromosome missegregation. Chromosome missegregation produces aneuploid karyotypes, a hallmark of cancer^27^, and drives distinct gene expression changes^28,29^ that have been linked to DNA damage from chromosomes caught in the cytokinetic cleavage furrow^30^, altered nuclear morphology^31^, micronucleus formation and rupture^32^, and gene dosage imbalances^33,34^. Which of these features are causally related, however, remains difficult to establish, because cells that missegregate chromosomes are highly heterogeneous, both in terms of their karyotypes and nuclear morphology (Fig. 4a); resolving the relationships therefore requires morphology, karyotype and gene expression to be measured in the same cell, an ability that is provided by the Waldo method.

**Fig. 4.**
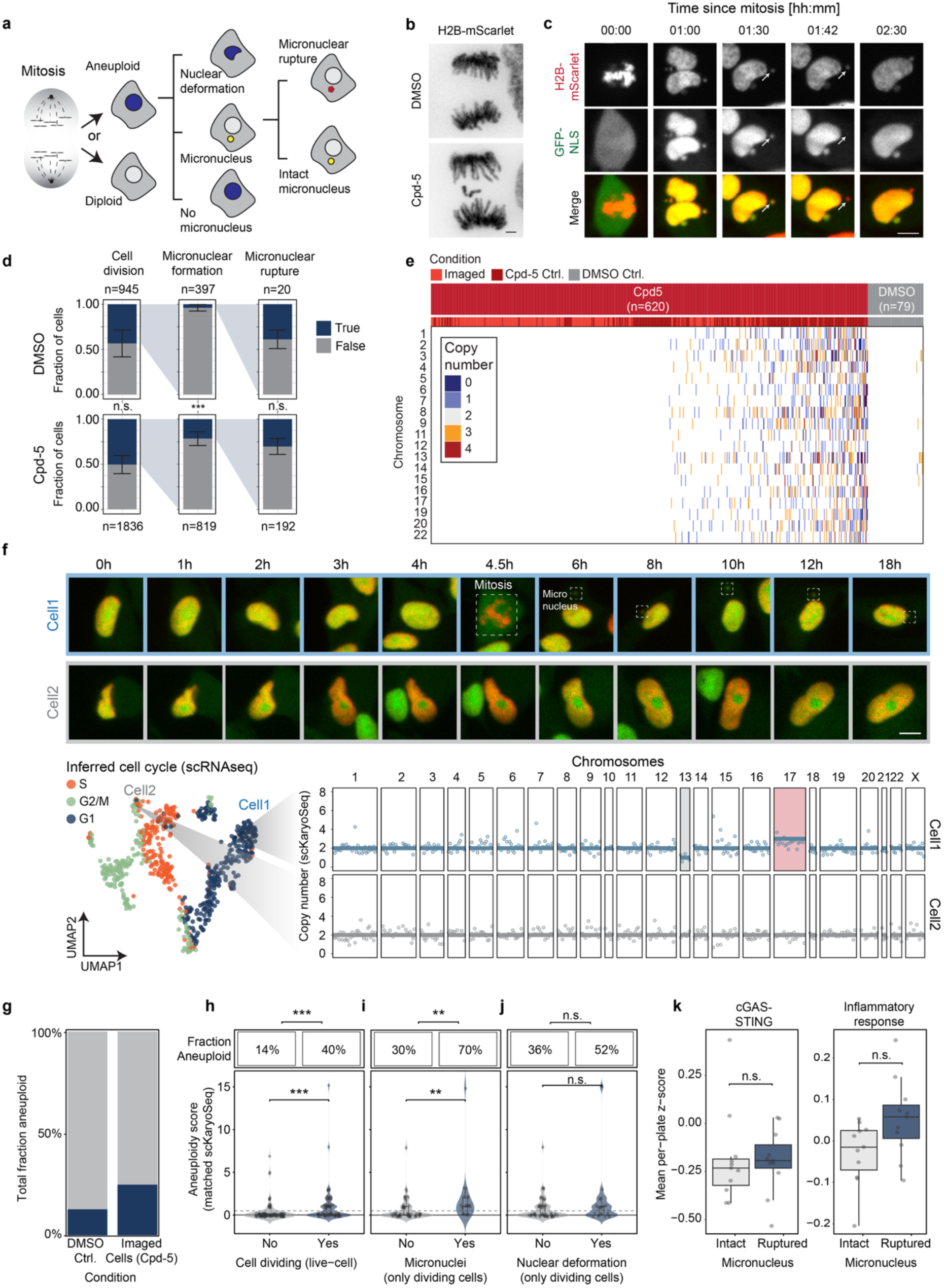
Studying immediate cellular responses to aneuploidy using Waldo. **a**, Cellular fates after mitosis: daughter cells after mitosis either contain a normal or abnormal karyotype. Cells with an abnormal karyotype may show nuclear deformation or form a micronucleus. Micronuclei may subsequently rupture or remain intact. **b**, Chromosome segregation (H2B-mScarlet) under DMSO or the Mps1 inhibitor Cpd-5. A lagging chromosome can be observed in a Cpd-5-treated cell. **c**, Timelapse of a dividing cell producing multiple micronuclei, one of which ruptures (arrow). Micronucleus rupture is observed as a sudden loss of GFP-NLS signal, while H2B-mScarlet signal is retained. **d**, Fraction of cells positive (True, dark blue) or negative (False, grey) for cell division, micronuclear formation and micronuclear rupture, under DMSO (top) and Cpd-5 treatment (bottom). Bars show the mean fraction across experiments, error bars the s.d.; n, total cells per phenotype. Significance by Cochran–Mantel–Haenszel test stratified by experiment with Benjamini–Hochberg correction: micronuclear formation increased under Cpd-5 (P_adj_=1.2×10^−12^), whereas cell division frequency (P_adj_=0.073) and micronuclear rupture (P_adj_=0.45) did not differ. n.s., not significant. **e**, Single-cell chromosome copy-number profiles (scKaryoSeq) for Cpd-5 (n=620) and DMSO (n=79) treated cells passing quality filters. Each column, one cell; each row, a chromosome; colour, inferred copy number (0–4). The top bars mark the sample type: Cpd-5 treated cells (red) for either cells that were imaged (light red) or grown in a separate well that was not imaged, used as a positive control (dark red) and DMSO treated control (grey) cells. Chromosomes 10, 18, 21 and X were excluded from the analysis, as data was too noisy (See Methods). **f**, Multimodal readout of two example cells. Top, live-cell imaging over 18 h (H2B-mScarlet, red; GFP-NLS, green); Cell1 divides and forms a micronucleus (dashed box), Cell2 does not divide. Bottom left, Harmony-integrated scRNA-seq UMAP colored by inferred cell-cycle phase (S, G2/M, G1; Seurat CellCycleScoring), with Cell1 and Cell2 indicated. Bottom right, scKaryoSeq profiles of both example cells. Points, normalized counts per genomic bin; line, whole-chromosome copy number. Missegregated chromosomes are shaded (red, gain; grey, loss). **g**, Fractions of aneuploid cells according to scKaryoSeq for each treatment condition (same as in **e**). Note that the Cpd-5 treated cells that were imaged show a lower fraction of aneuploidy cells compared to non-imaged cells, which is likely due to a variation in Cpd-5 drug concentration and possibly due to partial light-induced cell cycle arrest, especially at later time-points. **h–j**, Imaging phenotypes compared with matched single-cell karyotypes. Each cell is a matched imaging–sequencing pair. Upper boxes, fraction aneuploid (Fisher’s exact test on aneuploid vs. diploid for each phenotype); lower violin-and-box plots, per-cell distance from diploid karyotype (matched scKaryoSeq) for phenotype-negative (No) and -positive (Yes) cells (two-sided Wilcoxon rank-sum test); boxes, median and IQR. The dashed horizontal line separates diploid cells (below) from aneuploid cells (above). **h**, Fraction aneuploid cells for dividing versus non-dividing cells (n=131,78; P=8.5×10^−6^). **i**, Among dividing cells, fraction of aneuploid cells for cells with or without micronucleus (n=57,20; P=2.7×10^−3^). **j**, Among dividing cells, fraction of aneuploid cells for cells with or without nuclear deformation (n=56,21; P=0.20; n.s.). **k**, Pathway scores (mean per-plate z-score) in micronucleus-positive cells with an intact versus ruptured micronucleus, for curated cGAS–STING (left) and Hallmark inflammatory-response (right) gene sets. Each point, one cell (n=11 intact, 11 ruptured); boxes, median and IQR; whiskers, 1.5× IQR. Two-sided Wilcoxon rank-sum test with Benjamini– Hochberg correction across the two pathways; neither significant after correction (inflammatory response n.s.; cGAS–STING n.s.). Scale bars, 2 µm (b), 10 µm (c, f).

To this end, we applied Waldo to near-diploid RPE-1 cells transduced with a library of ten Waldo receptors. Waldo receptors showed the expected random, combinatorial expression patterns required for Waldo barcoding (Suppl. Fig. 4; Methods). Waldo RPE-1 cells also expressed H2B-mScarlet to follow chromosome segregation and micronucleus formation, and GFP-NLS to detect micronucleus rupture through loss of the GFP-NLS signal from micronuclei upon rupture (Fig. 4b,c, Suppl. Video 2, Methods). Chromosome missegregation was induced using Cpd-5, an inhibitor of the spindle assembly checkpoint kinase Mps1^35^, and cells were tracked by time-lapse spinning disk confocal microscopy for 20–24 h. Fluorescent barcode readout was performed during the last 2-3 h of time-lapse imaging, with rounds of barcode readout being performed in between live-cell imaging time-points (Suppl. Video 3, Methods). Barcode readout did not compromise cell retention (Suppl. Fig. 5a) or cause severe cellular stress, as cells continued to enter mitosis throughout the readout rounds (Suppl. Fig. 5b).

For phenotypic single-cell readouts, we first analyzed cell division in live-cell imaging to identify cells with micronucleus formation and rupture, and nuclear deformation after division. Cpd-5 treatment did not alter the number of cells undergoing cell division—approximately 50% of cells underwent cell division during the experiment in both conditions—but increased the fraction of cells showing micronuclei formation post-division (3.9% in DMSO vs. 21.6% in Cpd-5), a subset of which subsequently ruptured (∼35% in both conditions, Fig. 4d).

After imaging, cells were photoactivated, collected by FACS and processed for scRNA-seq to obtain transcriptomic information and scKaryoSeq^36^ to determine karyotypes of single cells (Fig. 4a; Methods). Of photoactivated cells, ∼40% were successfully recovered by FACS sorting, of which ∼28% passed quality control filters for single-cell sequencing (Suppl. Fig. 5c). Both sequencing library depth and the fraction of cells passing quality control were comparable between imaged cells and non-imaged control cells plated in parallel (Suppl. Fig. 5d,e).

Both scKaryoSeq and scRNA-seq can be performed on Waldo-barcoded cells without modification of the Waldo workflow; cell-specific barcodes are appended to the DNA probes following EagI restriction digestion and ligation of sample adapters, and because EagI also cuts genomic DNA, adapter-ligated genomic fragments—which report on chromosome copy number—can be co-captured alongside cellular mRNA during library preparation (Suppl. Fig. 3a). To confirm that combining scKaryoSeq and scRNA-seq during sample preparation does not compromise either method, we first applied the Waldo workflow to A549 cells, an aneuploid cancer cell line, as a control. Total unique mRNA counts per cell were comparable between the Waldo and standard CEL-Seq2 protocols (Suppl. Fig. 6a), and karyotype measurements closely resembled those from standard scKaryoSeq, with both protocols detecting the known amplification of the left arm of chromosome 2 (Suppl. Fig. 6b,c) ^37^.

Having established a workflow for combining scKaryoSeq and scRNA-seq, we next processed the RPE-1 Waldo samples that were analyzed by time-lapse imaging. Analysis of single-cell karyotype sequencing confirmed that Cpd-5-treated cells showed increased aneuploidy, with both increases in chromosome copy-number gains and losses compared to untreated cells (Fig. 4e).

Next, the sequencing-based barcode was matched to the corresponding imaging-based barcode using a two-stage algorithm. In the first stage, a soft Jaccard score compared which receptors were expressed in each cell to narrow the pool of imaged cells down to a small group of candidates that represent most likely matches. In the second stage, the best match within the candidate pool was selected by Pearson correlation analysis across the expression levels of all ten receptors (Methods). Barcode matching yielded a total of 231 matched cells between imaging and sequencing. Two example cells are shown in Fig. 4f, revealing their live-cell imaging history, inferred cell cycle phase at the end of imaging (from scRNA-seq) and scKaryoSeq chromosome copy-number profile. Live-cell imaging showed that Cell 1 underwent cell division and formed a micronucleus, suggesting that Cell 1 had missegregated one or more chromosomes. Further analysis of live-cell imaging data showed that the micronucleus had not ruptured when cells were harvested. Karyotype analysis of the same cell confirmed one monosomy (chromosome 13) and one trisomy (chromosome 17), consistent with the missegregation event inferred from imaging. In contrast, Cell 2 did not divide during the imaging experiment and, therefore remained diploid as determined by karyotype analysis (Fig. 4f).

The example cells described above not only illustrate the power of the Waldo method to study cell division, but also implicitly suggest that cell linking between imaging and sequencing modalities was correct, since the dividing, but not the non-dividing example cell was matched with an aneuploid cell in the sequencing dataset. To validate the accuracy of barcode matching more quantitatively, we therefore leveraged the fact that ∼50% of cells did not undergo cell division during the experiment and should therefore show background aneuploidy rates (∼12%, see Fig. 4e,g). In contrast, cells that did undergo cell division in the presence of Cpd-5 should have increased levels of aneuploidy. Based on the background level of aneuploidy, the total fraction of cells that was aneuploid in the imaged cells in Cpd-5 treatment and the fraction of cells that underwent division, we calculated that ∼38% of dividing cells should contain an aneuploid karyotype (Methods). After cell matching between imaging and sequencing, we found that 14% of non-dividing and 40% of dividing cells were aneuploid (Fig. 4h), indicating that barcode matching linked the majority of cells correctly.

Having established that barcode matching is accurate for the majority of cells, we next asked how aneuploidy relates to nuclear morphology. Among dividing cells, cells that formed a micronucleus were significantly more aneuploid than cells that did not (Fig. 4i; 30% vs 70% aneuploidy for cells without or with a micronucleus, respectively), providing a direct comparison of the degree of aneuploidy accompanying micronucleus formation. Earlier studies have reported that nuclear deformation is also associated with aneuploidy^31,38,39^. In our data, however, we did not observe a statistically significant association between nuclear deformation and aneuploidy (p = 0.2), although a trend towards higher levels of aneuploidy in cells with nuclear deformation post-division was observed (52% vs 36% aneuploidy for cells with and without nuclear deformation, respectively) (Fig. 4j). Nuclear deformation is therefore not associated with aneuploidy at the levels induced here, suggesting that the association reported previously^31^ may require more severe karyotype changes. Together, the comparisons above show that linking live-cell imaging to scKaryoSeq resolves which post-mitotic morphological features are associated with aneuploidy.

Finally, we asked whether micronucleus rupture alters gene expression, focusing on inflammatory signaling and specifically on the cGAS-STING pathway, which has been proposed to be triggered when micronucleus rupture exposes DNA to the cytoplasm^40,41^. Cells with a ruptured micronucleus showed a modestly elevated inflammatory-response transcriptomic signature relative to cells with an intact micronucleus. However, after correction for multiple testing, the comparison did not reach significance (Fig. 4k). A curated cGAS-STING signature was similar between the two groups (Fig. 4k, Suppl. Table 3). At the early post-division time-point captured here, micronucleus rupture is therefore associated with at most a modest elevation of the inflammatory transcriptional response, consistent with recent findings using bulk RNA sequencing of cells with or without micronuclear rupture^42^. Together, these analyses connect the morphological consequences of chromosome missegregation to karyotypes and transcriptional states in individual cells. More broadly, they establish Waldo as a general approach for linking imaging-based phenotypes to single-cell sequencing readouts in the same cell.

## Discussion

We have developed Waldo, a genetically encoded single-cell barcoding technology whose barcodes can be read out by both live-cell imaging and single-cell sequencing. To our knowledge, Waldo is the first barcode that can be measured directly in both modalities, and Waldo therefore provides a general approach to link a cell observed by imaging to the same cell profiled by sequencing. Because the barcode is displayed at the cell surface and read out without detaching or isolating cells, Waldo is compatible with high-resolution live-cell imaging of cells in their native context, and we showed that Waldo integrates with at least two single-cell sequencing readouts without compromising sequencing quality. A key feature of Waldo is the iterative fluorescence readout, in which probes are stained, imaged and then removed by a nuclease before the next round. By reusing the same fluorescence channel across successive rounds, iterative readout decouples barcode diversity from the number of spectrally distinct fluorophores available, making the achievable barcode diversity, in principle, unlimited. Our simulations indicate that barcode diversity scales with the number of receptors and is substantially improved by reading out expression level rather than the mere presence or absence of each receptor. Barcode space can therefore be expanded either by adding receptors or by broadening the range of expression levels.

Several developments could extend Waldo further. Since Waldo barcodes can be read out both in transcriptomes and by PCR amplification of DNA probes, future protocols could combine Waldo with established single-cell sequencing approaches for profiling protein–DNA interactions^26,43^, ribosome profiling^44^, RNA stability^45^ and chromatin marks^24,46,47^. As such, Waldo could become applicable to a broad set of biological questions linking cellular phenotypes to different aspects of gene expression control. Additionally, to increase barcode diversity further, advances in de novo peptide binder design should make it straightforward to expand the Waldo receptor-probe set to much larger complexities^48,49^. Combining a genetic barcode such as Waldo with spatially resolved molecular barcoding approaches, including light-directed oligonucleotide barcoding^50,51,52^ and array- or microfluidics-based spatial oligonucleotide barcoding strategies^13,14,15,16^, could further increase barcode diversity. Combining Waldo with targeted photoactivation-based enrichment of cells with rare imaging-based phenotypes would allow analysis of larger numbers of such cells to understand the origin of rare phenotypes. Isolation of rare cells through photoactivation is already applied in various methods^42,53^, and can further build on the dye-based photoactivation approach developed here. Finally, because the Waldo barcode is encoded by expressed receptor proteins, the barcode also exists at the protein level and could serve as a single-cell protein-based barcode^54^. As single-cell mass spectrometry matures, protein-level detection of barcodes could allow Waldo to bridge imaging to single-cell proteomics.

While Waldo functions well with the current set of reagents, the current system does have several limitations. First, barcodes are introduced by lentivirus, so only model systems that are permissive for lentiviral transduction are accessible to the Waldo approach. Second, the barcode is genetically encoded and therefore clonal: every cell derived from a barcoded founder cell inherits the same barcode, so clonally related cells cannot be distinguished from one another by Waldo. In cell lines, seeding the imaged cells from a large, diverse starting population makes it unlikely that any two clonal relatives are present in the same experiment, ensuring that each barcode appears only once among the imaged cells. In model organisms, however, where cells cannot be dissociated and reseeded, clonal relatives are unavoidable. An inducible cassette that drives random expression of a receptor subset through Cre/lox-mediated recombination (e.g. Brainbow^18^) could address both limitations by removing the need for lentiviral transduction, providing temporal control over when barcodes are induced, and enabling unique barcode generation within clonal populations^21^.

A related challenge is that sister cells arising from a division during the experiment also carry identical barcodes. In our proof-of-concept experiments we limited the problem by seeding cells shortly before imaging and allowing at most one division per cell. Having two sisters within the cell population may not always be detrimental, as in many cases sister cells are similar both in terms of genotype and phenotype, so mis-assigning them may not have a major impact on data interpretation. However, in some cases discriminating sisters is critical, for example when a division is asymmetric—one daughter forms a micronucleus while the other does not. In such cases, matching cells to the wrong sister adds noise to analyses, requiring a larger number of cells for statistical power. Photoactivating or photo-crosslinking oligonucleotides to only one of the two sisters would resolve the ambiguity, and recent advances in on-the-fly automated image analysis and photoactivation should make targeted photoactivation feasible^42,53^.

A third limitation is the barcode diversity of the current receptor set. With the ten receptors used in this study, the number of cells profiled per experiment (200–400) approached the upper bound of the achievable barcode diversity, such that some cells carried barcodes that were very similar to multiple other cells, which may have resulted in a minority of cells being matched incorrectly. Adding further receptors, or applying the strategies outlined above, should readily increase barcode diversity above the cell numbers profiled and allow the approach to scale to larger experiments with few matching errors. Since completing the proof-of-concept experiments, we have already developed two additional receptor–probe pairs, which will be made available upon publication.

Live-cell imaging and single-cell sequencing have developed largely independently, the first resolving cellular behavior over time, the second the molecular state of individual cells. Waldo introduces a single barcode that can be read out by both, allowing the two measurements to be made for the same cell and related to one another. We anticipate that connecting dynamic phenotypes to genome-wide molecular state will make a broad range of questions in cell biology accessible that neither approach can address alone.

---

**Supplementary Fig. 1.**
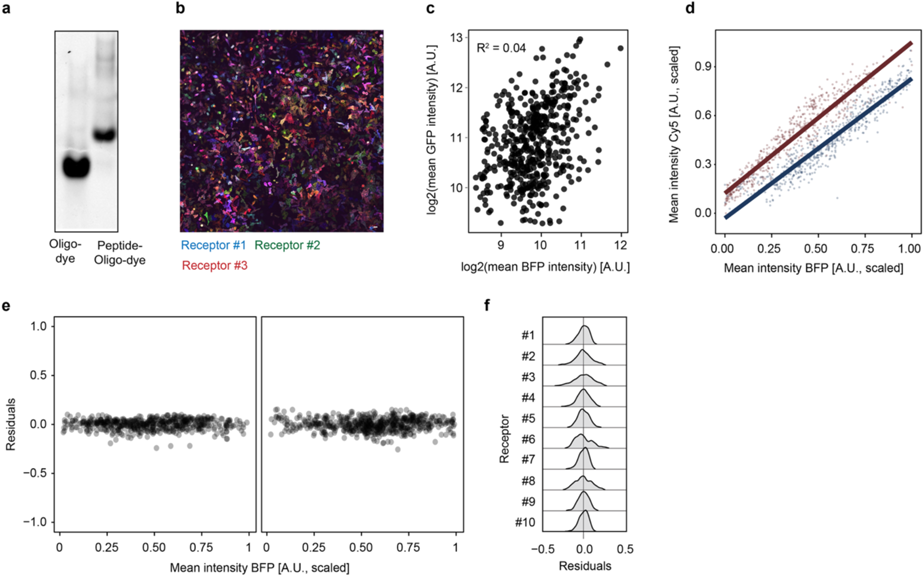
Characterization of receptor expression and quantitative readout of Waldo barcodes. **a**, In-gel fluorescence of the Cy5-labelled oligo before (left) and after (right) conjugation to the peptide. **b**, Representative image of HeLa cells co-transduced with three Waldo receptors, each marked by a distinct fluorescent protein. Receptor #1 (BFP, blue), Receptor #2 (GFP, green) and Receptor #3 (mCherry, red). Various color combinations across single cells indicate independent transduction of receptors. **c**, Quantification of the cells in **b**, showing single-cell Receptor #1 (BFP) versus Receptor #2 (GFP) expression levels. Each point represents one cell (n=484). The low R^2^ (0.04) indicates that the two receptors are expressed independently. **d**, Mean probe (Cy5) intensity versus mean receptor (BFP) intensity for two randomly chosen receptors (dark red and dark blue), with a linear fit for each receptor. Probe signal scales linearly with receptor expression. **e**, Residuals of the linear fits in **d** plotted against receptor (BFP) intensity. Residuals are evenly spread across the intensity range, indicating a linear and homoscedastic probe readout. **f**, Distribution of fit residuals for each of the ten receptors (Receptor #1–#10). Residuals are approximately Gaussian. Scale bar, 20 µm (b).

**Supplementary Fig. 2.**
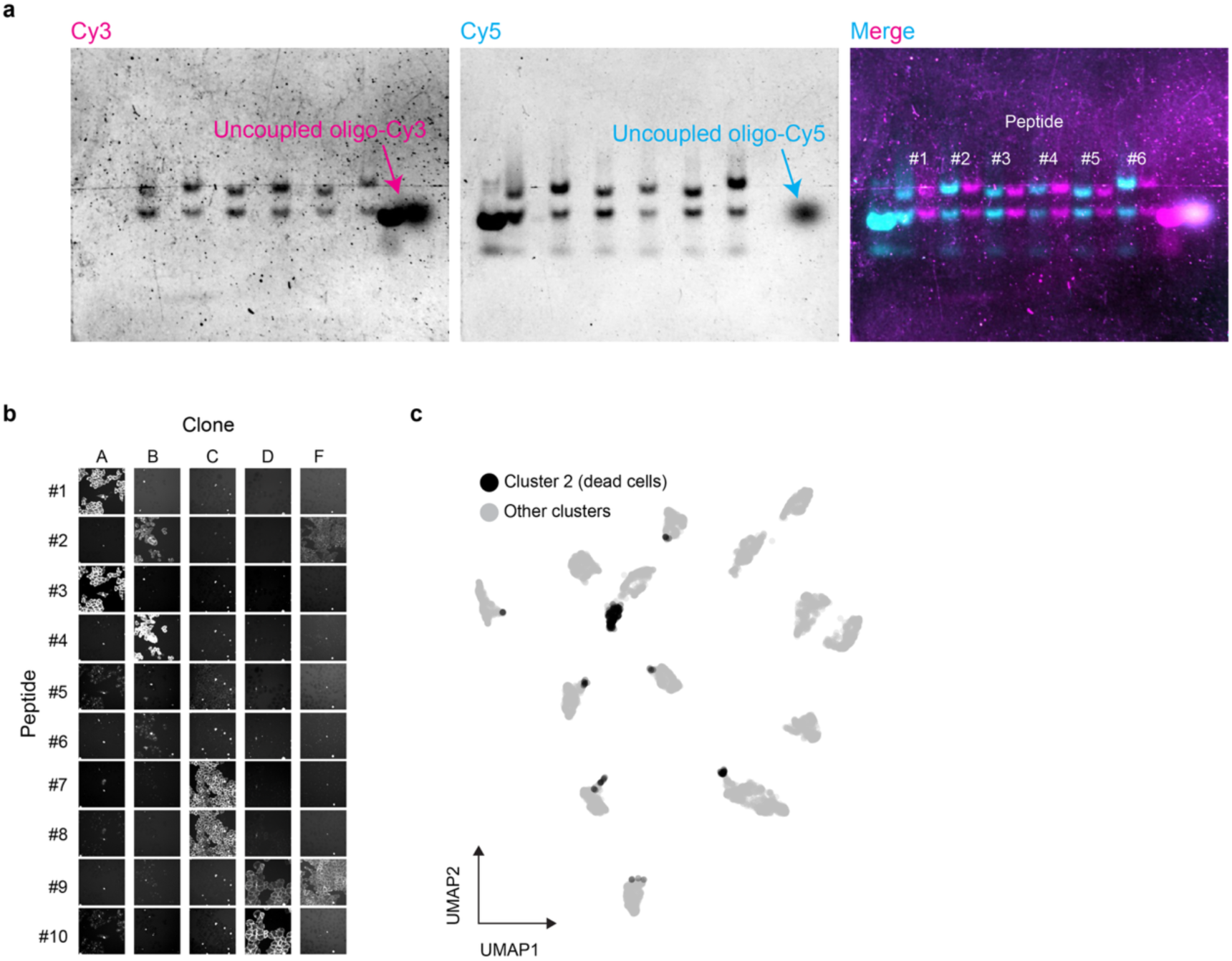
Probe conjugation and fluorescent barcode readout of clones. **a**, In-gel fluorescence of the first six peptide probes (Peptide #1–#6), each conjugated to a Cy3-labelled and a Cy5-labelled oligo and imaged in two channels. Left, Cy3 channel. Middle, Cy5 channel. Right, merge (Cy3, magenta; Cy5, cyan). In each lane the upper band is the conjugated peptide–oligo–dye product and the lower band the unconjugated oligo–dye. Arrows mark control lanes containing uncoupled oligo-Cy3 and oligo-Cy5. Coupling efficiency varies slightly among different probes. **b**, Representative probe staining images of five barcoded monoclonal cell lines (clones A, B, C, D and F) across ten probes (Probe #1–#10). Each clone shows a distinct staining pattern. **c**, UMAP of single-cell barcode intensities (as in Fig. 2g), with Phenograph cluster 2 highlighted in black. This cluster corresponds to dead cells.

**Supplementary Fig. 3.**
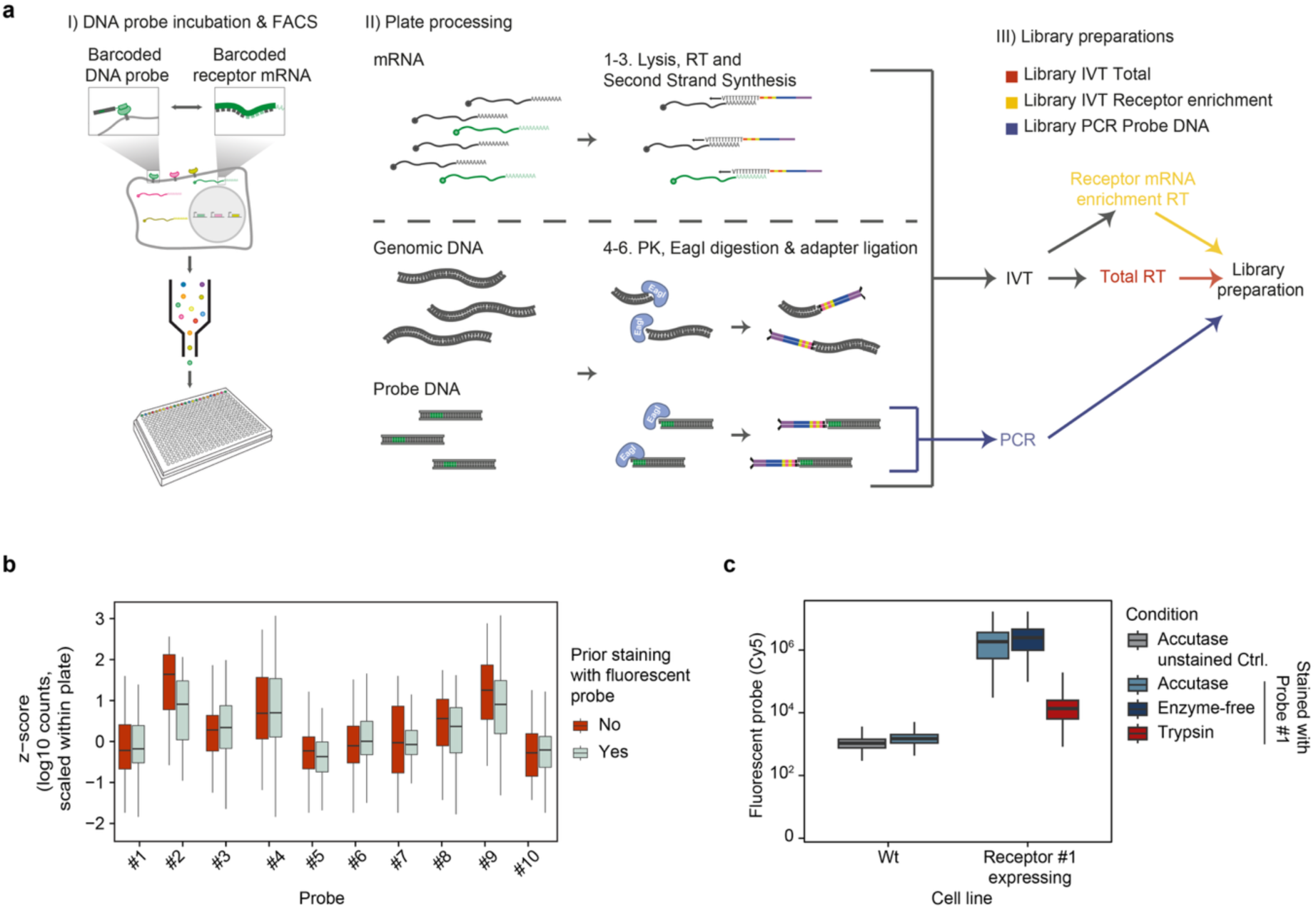
Single-cell sequencing workflow, imaging compatibility and receptor integrity after cell detachment. **a**, Schematic of the single-cell sequencing readout. I) Cells are stained with barcoded DNA probes and sorted by FACS into plates. II) In each cell, mRNA is reverse-transcribed and converted to double-stranded cDNA using the CEL-Seq2 protocol. Subsequently, genomic DNA and DNA probes are digested with EagI and ligated to sample adapters. III) Cells are pooled and then split into two samples for preparation of three different libraries. DNA probes are amplified by targeted PCR (Library PCR Probe DNA). The remainder underwent linearly amplification by *in vitro* transcription (IVT) following the scDam&T approach^55^, after which reverse-transcription was performed with either random primers (Library IVT Total) or receptor-targeted primers that enrich receptor mRNAs (Library IVT Receptor enrichment). **b**, Sequencing read counts per cell for each DNA probe, shown as z-scores of log_10_-transformed counts and scaled within each sequencing plate. Cells were either imaged and subjected to fluorescent barcode readout or grown in a separate well that was not imaged (No imaging 1 and 2). Boxes show median and IQR, whiskers 1.5× IQR (outliers not shown). **c**, Effect of the cell detachment reagent on receptor integrity, as determined by fluorescent probe staining intensity by flow cytometry. Left, wild-type cells without receptor expression detached with Accutase, either unstained or stained with Probe #1. Right, Receptor #1-expressing cells detached with Accutase, enzyme-free solution or trypsin and stained with Probe #1. Trypsin reduces staining ∼100-fold compared with Accutase or enzyme-free detachment. Boxes show median and IQR, whiskers 1.5× IQR. n=28,500 cells per condition.

**Supplementary Fig. 4.**
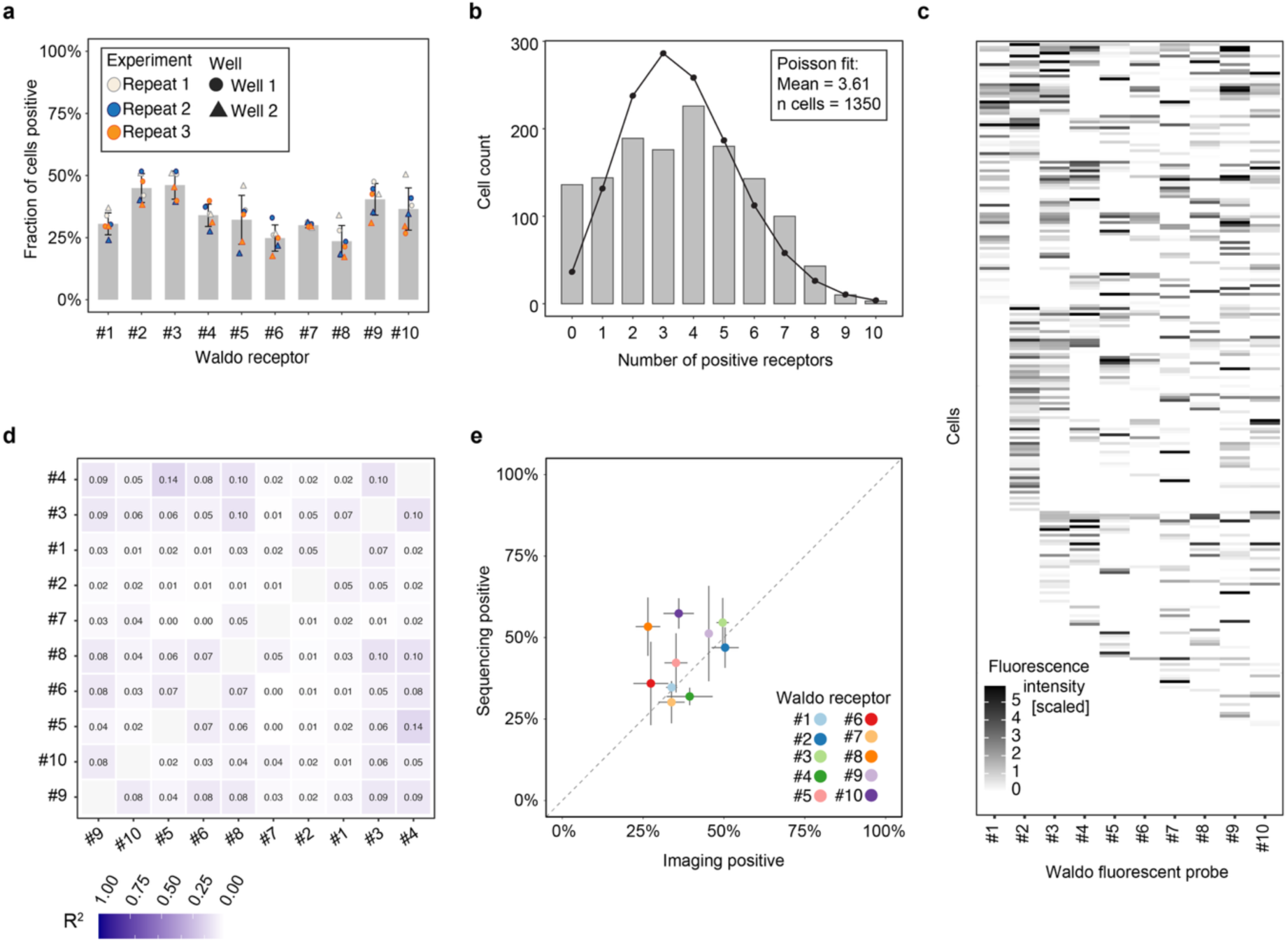
Characterisation of barcode expression in the RPE-1 Waldo cell line. **a**, Fraction of cells scoring positive in imaging for each of the ten fluorescent probes. Bars show the mean and error bars the s.d. across conditions. Individual points show each group, coloured by experiment (Repeat 1–3, three biological repeats) and shaped by well (circle, Well 1; triangle, Well 2). **b**, Distribution of the number of receptors expressed in single cells (Well 1, pooled across the three experiments; n=1,350 cells), overlaid with a maximum-likelihood Poisson fit (mean=3.61). **c**, MAD-scaled probe (Cy5) intensity for the ten fluorescent probes (columns) across single cells (rows). Cells are sorted by their binary barcode string. **d**, Pairwise R^2^ between the probe (Cy5) intensities of all fluorescent probes measured by imaging. Probe order follows hierarchical clustering. Low R^2^ values indicate that receptors are expressed independently. **e**, Fraction of cells positive for each receptor by imaging versus by sequencing, each averaged across the three experiments (Well 1). Each point is one Waldo receptor. Horizontal and vertical lines show the s.d. across experiments for imaging and sequencing, respectively. The dashed line indicates perfect agreement between imaging and sequencing.

**Supplementary Fig. 5.**
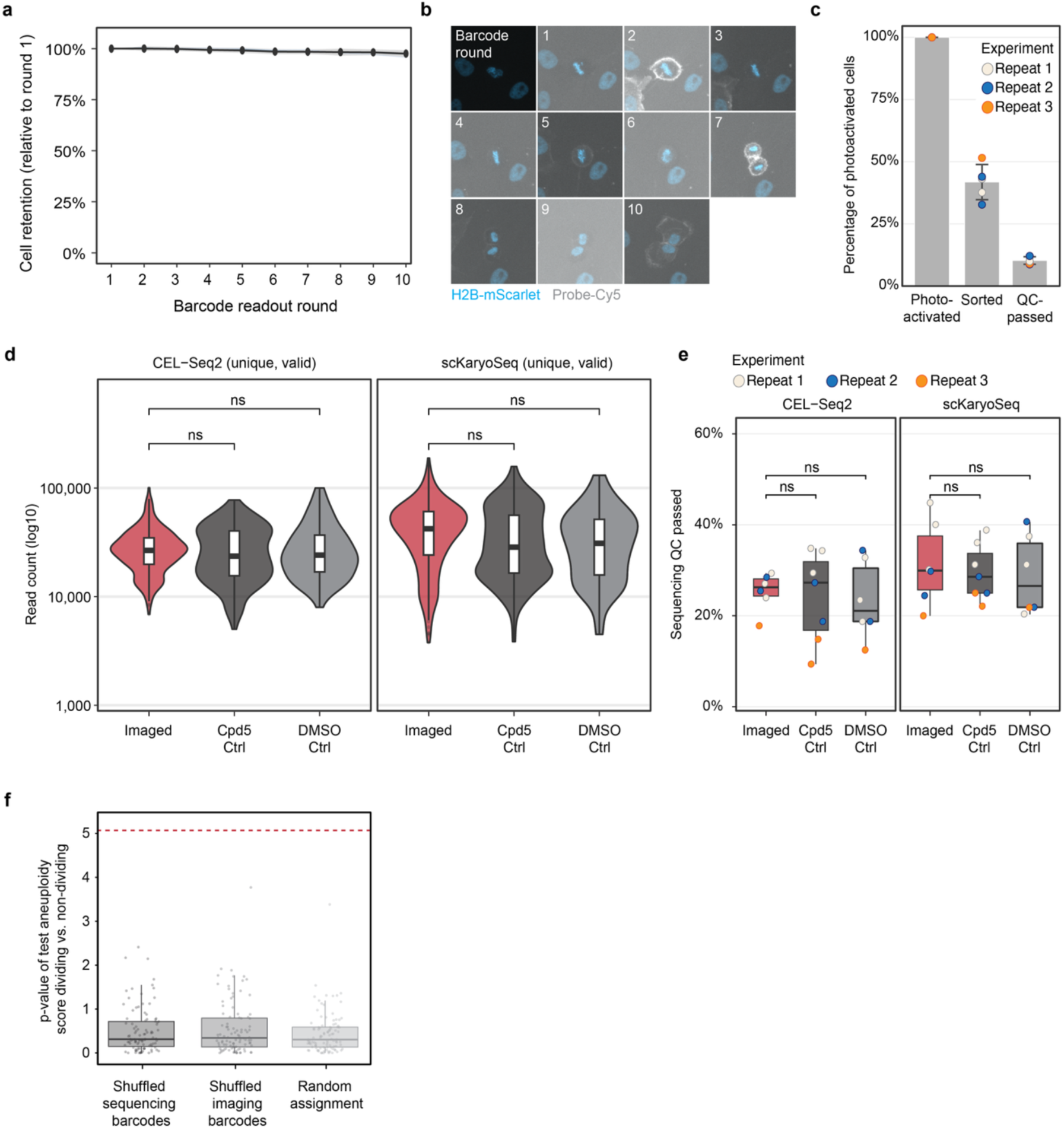
The Waldo barcode readout preserves cell retention and does not compromise single-cell sequencing quality. **a**, Cell retention on the plate across the ten sequential barcode readout rounds, expressed as the fraction of segmented cells detected relative to round 1. The line shows the mean and the shaded band the s.d.. Values near 100% indicate negligible cell loss. **b**, Representative images across the ten barcode readout rounds for a single cell that divides during the protocol, showing that cells remain attached and continue to cycle throughout sequential barcode readout rounds. Cyan, H2B-mScarlet. Grey, fluorescent probe (Probe-Cy5). **c**, Cell recovery at successive stages of the pipeline. Bars show the mean and error bars the s.d. of the percentage of photoactivated cells retained at each stage: photoactivation (100% by definition), FACS sorting and sequencing quality control. Individual points show each well (n = 5 wells), coloured by experiment (Repeat 1–3, three biological repeats). **d**, Sequencing library depth for cells passing quality control filters for each condition (Imaged, Cpd-5 control, DMSO control) and each modality (CEL-Seq2 and scKaryoSeq). Violin plots show the full distribution and embedded boxes show median and IQR. Imaged cells were compared with each control by two-sided Wilcoxon rank-sum test. n.s., not significant. **e**, Fraction of cells that passed sequencing quality control filters per plate for each condition and modality. Boxes show median and IQR. Individual points show per-plate values, colored by experiment (Repeat 1–3). Comparisons as in **d**. n.s., not significant. **f**, Controls for the barcode matching validation. Barcodes were shuffled and single cells were re-linked between imaging and sequencing modalities with new shuffled barcodes. The analysis of the aneuploidy score for dividing and non-dividing cells was performed as in Fig. 4. Statistical tests were then performed to determine whether dividing cells were more aneuploid than non-dividing cells. Boxes show the distribution of Mann–Whitney −log_10_(*P*) values across 100 permutation runs of barcode shuffling for each control condition. In shuffled sequencing barcodes, receptors were shuffled per cell across the sequencing barcode matrix. In shuffled imaging barcodes, receptors were shuffled per cell across the imaging fluorescence matrix. In random assignment, each sequencing cell was assigned to a randomly drawn imaging cell without using barcode profiles. Individual points show each permutation run, pooled across all six imaged wells. The dashed red line indicates the value obtained with the real barcode-matched pairs (−log_10_(*P*)=5.07, n=209 cells with both live-cell cell division annotation and karyotype data. All three controls produce values near zero.

**Supplementary Fig. 6.**
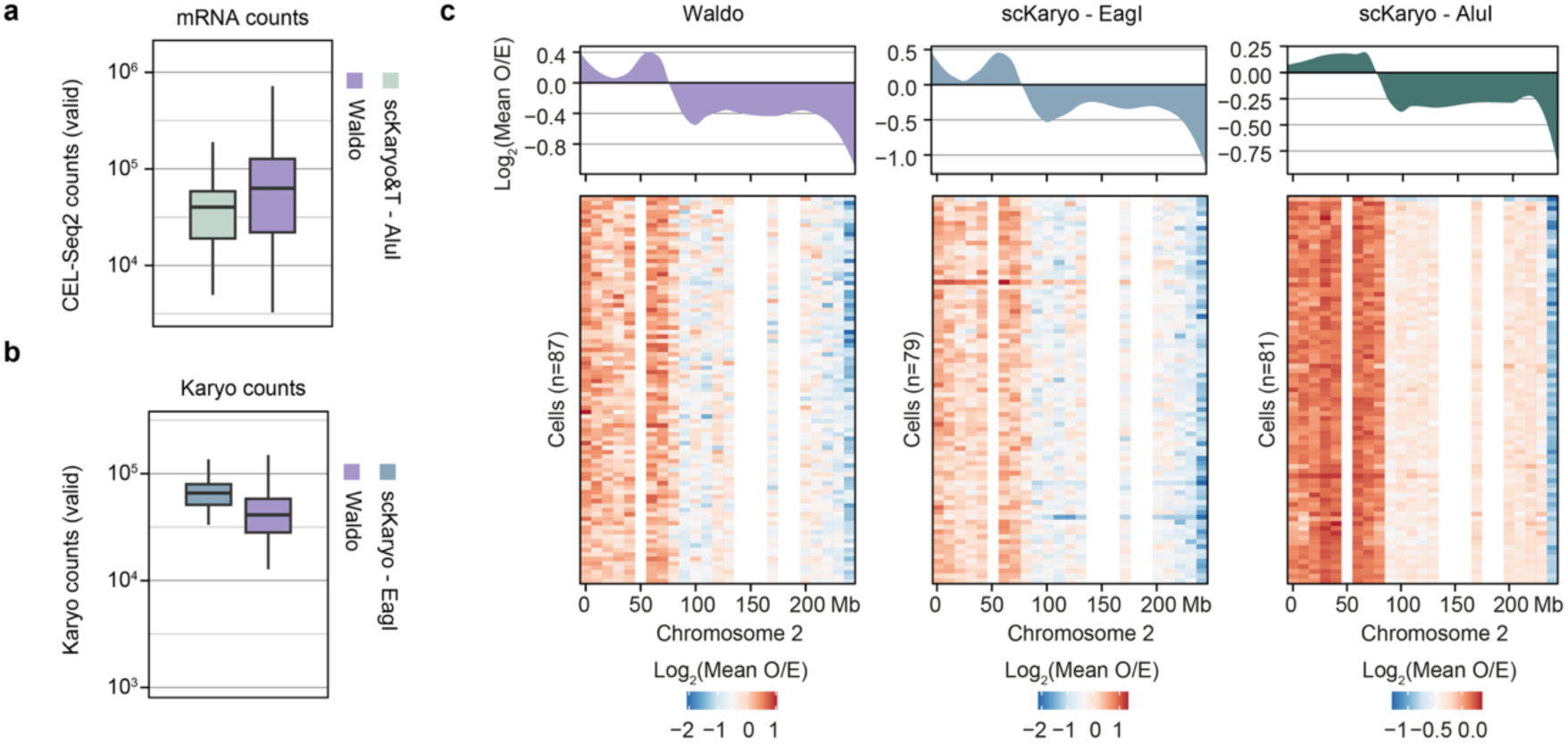
The Waldo protocol is compatible with single-cell mRNA and karyotype sequencing. **a**, Comparison of valid CEL-Seq2 counts per cell, a measure of transcriptome capture, for the Waldo and scKaryo&T–AluI (gold standard) datasets. n=283 (Waldo) and 77 (scKaryo&T–AluI) cells. **b**, Valid karyotype sequencing counts per cell for the Waldo and scKaryo–EagI (gold standard) datasets. n=308 (Waldo) and 88 (scKaryo–EagI) cells. For **a** and **b**, boxes show median and IQR, whiskers 1.5× IQR, points outliers. **c**, Copy-number profiles across chromosome 2 for the Waldo, scKaryo–EagI and scKaryo–AluI datasets. Top, pseudobulk profile (mean across all cells). Bottom, heatmap in which each row is one cell. Values are log_2_(mean observed/expected) in 10-Mb bins. n=87 (Waldo), 79 (scKaryo–EagI) and 81 (scKaryo–AluI) cells, each from a single representative plate. Note that the color scale and the axis range of the top panel differ between the three datasets.

## LLM disclosure

During the preparation of this work, the authors used ChatGPT (OpenAI) and Claude (Anthropic) to assist with language editing and code annotation and generation. Following their use, the authors thoroughly reviewed and revised all generated outputs and assume full responsibility for the content of the published article. The scientific concepts, analyses, interpretations, and conclusions were entirely devised, implemented, and verified by the authors.

## Code availability

All code relevant to this manuscript is deposited on github and will be made publicly available upon publication of the manuscript. For completeness during the revision process, a static snapshot of the current code has been uploaded to figshare https://figshare.com/s/91ec7e969427a29ab55d.

## Data availability

A selection of raw images that support the findings of this study are openly available at https://figshare.com/s/b90b0ee569822d701728 (Application experiments shown in Fig. 4, Suppl. Fig. 4-5) and https://figshare.com/s/b1d65bb88c88fe10703b (Receptor-probe specificity images and quantification shown in Fig. 1). All sequencing data from the application experiments (Fig. 4, Suppl. Fig. 4-5) have been deposited at the Gene Expression Omnibus (GEO) under accession number GSE338511.

## Author contributions

M.M., J.K. and M.E.T. conceptualized the study. M.M., S.J.A.L., L.A., R.B., J.F.M. and C.D.M. performed the experimental investigation. M.M., P.M.J.R., M.M.G., B.A.K. and V.P. conducted formal analysis. M.M., J.K. and M.E.T. wrote the original draft of the manuscript. All authors reviewed and edited the manuscript. J.K. and M.E.T. supervised the study and acquired funding.

## Acknowledgments

We thank members of the Tanenbaum lab for helpful discussions. Clément Rouillon, Roxanne van der Sluijs, Charlotte Delvaux de Fenffe and Francesca Mattiroli for help with running gels. Sandra de Vries for her help and input on sequencing library preparation. M.M., G.J.P.L.K, J.K. and M.E.T. were supported by the Oncode Institute, which is partly funded by the Dutch Cancer Society (KWF). M.M. and M.E.T. were further supported by a grant from the European Union (ERC, VirIm, 101044794).

## Methods

### Experimental methods

#### Cell culture

Cell lines were maintained in a humidified chamber at 37 °C in 5% CO2 and were routinely screened for mycoplasma. A549, RPE1, U2OS and HT1080 cells (A549 - ATCC, CCL-185; RPE1 – ATCC, CRL-4000; U2OS - ATCC, HTB-96; HT1080 – from Kind et al. 2013^56^) were grown in high-glucose Dulbecco’s Modified Eagle Medium (DMEM) containing GlutaMAX (Gibco, 31966021), and supplemented with 10% FBS (Sigma, F7524) and 1% Pen/Strep (Gibco, 15140122). HeLa and HEK293T cells (HeLa – ATCC, CCL-2; HEK293T – ATCC, CRL-3216) were grown in high-glucose DMEM containing GlutaMAX, supplemented with 5% FBS and 1% Pen/Strep. Cells were split every 2–4 d. For routine collection, cells were rinsed with Phosphate Buffered Saline (1X PBS, pH 7.4, in-house) and incubated with TrypLE Express Enzyme (Gibco, 12605010) for 5 min at 37 °C. Cells were dissociated by pipetting, and TrypLE was quenched by diluting the cells fivefold in culture medium before continuing with further sample processing as described in the next sections.

#### Plasmids

Lentiviral constructs were generated for each Waldo receptor using the pHR plasmid, which contains a SFFV promoter and WPRE element for high cellular expression. The Waldo receptor is composed of the following elements (5’ to 3’) – the CD40-ligand module (transmembrane domain), a fluorophore (e.g. BFP, mCherry or GFP), a GS-linker and a unique receptor for each construct. For experiments with RPE-1 cells, the pHR plasmid was used to generate constructs containing either sfGFP-NLS or H2B-mScarlet. All plasmids generated in this study will be deposited at and made available through Addgene.

#### Lentiviral production and cell line transduction

To produce lentivirus, HEK293T cells were transfected using Polyethylenimine (PEI, Polysciences Inc, 23966) with the lentiviral plasmid of interest and packaging vectors psPax and pMD2.G. The cell culture medium was refreshed 1 day after transfection. The supernatant containing the lentivirus was collected 3 days after transfection. The lentivirus was 100-fold concentrated using Lenti-X™ Concentrator (Takara Bio, 631232) and stored at –80°C in single-use aliquots. Before use, the titer of each lentivirus was determined. The different lentiviruses (each encoding a specific Waldo receptor) were diluted to equal titers and pooled to generate the combined Waldo lentiviral library. For generation of cell lines stably expressing the Waldo receptors, cells were incubated with lentivirus overnight together with 1ul of Polybrene (Merck, TR-1003-50UL) per 3ml of medium, after which the medium was replaced and cells were cultured for at least 4 days before further analysis was performed.

#### Generation of Waldo probes

##### Fluorescent probes

The fluorescent probe consists of a peptide (which binds to a specific Waldo receptor) synthesized with a N-terminal cysteine (GenScript) (extended with a GS linker to a total length of 25 amino acids), a single-stranded DNA (ssDNA) linker (30 nt, with a 5’ azide group (N3, IDT, /5Azide/); sequence in Supplementary Table 2) and a fluorophore (Cy3 or Cy5). Fluorescently labelled oligos were ordered from IDT. The probe was assembled in two steps. First, DBCO-PEG4-maleimide linker (Sigma, 760676; 2 nmol/µL in DMSO) was conjugated to an azide-labelled oligo by combining 20 nmol linker (10 µL) with 4 nmol oligo (40 µL) and incubating for 2 h on a rotor at room temperature. The oligo–linker was purified through an Amicon Ultra Centrifugal Filter, 3 kDa MWCO, washing at least three times (top up to 500 µL with 1X PBS, pH 7.4, 14,000 g, 10 min, room temperature) and recovered by inverting the filter into a fresh vial and spinning at 1,000 g for 2 min. Second, the oligo–linker was conjugated to the peptide (1 nmol/µL in 1X PBS, pH 7.4) via the maleimide–cysteine reaction by combining 8 nmol peptide (8 µL) with the oligo–linker and incubating for 1.5 h on a rotor at room temperature, followed by overnight incubation at 4 °C. The coupled probe was purified through an Amicon Ultra Centrifugal Filter, 10 kDa MWCO, using the same wash procedure. Finally, fluorescent probes were aliquoted and stored at −20 °C. Probe concentration was determined by NanoDrop™ 2000 and the molar concentration was calculated from the measured DNA concentration and the expected oligo length. To assess probe integrity and coupling efficiency, samples were normalized to 50 nM in 1X PBS, pH 7.4 and 5 µL was loaded onto a 6% polyacrylamide gel alongside a DNA ladder. Successful coupling was confirmed by the expected mobility shift of the peptide–oligo product relative to the free oligo.

##### DNA probes

The DNA probe consists of a peptide (recognizing a specific Waldo receptor) synthesized with a N-terminal cysteine (GenScript) (extended with a GS linker to a total length of 25 amino acids) and a double-stranded DNA adapter carrying a probe-specific barcode. All DNA probe and primer sequences as well as the universal linker sequence are included in Supplementary Table 2.

The DNA adapter part of the DNA probe was generated by PCR amplification of a universal linker sequence (pUC57 plasmid template, 300 bp). The unique forward primers (IDT) contain a 5’ phosphate group (/5Phos/), a 22 nt linker sequence with an EagI restriction-digestion motif, a 3 nt UMI, a 6 nt probe-specific barcode, a 3 nt UMI and a 24 nt primer binding sequence to anneal to the universal linker sequence. The common reverse primer (IDT) contains a 5’ azide (N3, /5Azide/) for downstream conjugation to the peptide and a 24 nt primer binding sequence to anneal to the universal linker sequence. Large-scale PCR amplification (2.5 mL total reaction volume) was performed with Q5 High-Fidelity DNA Polymerase (NEB, M0491S) following the standard protocol with 34 cycles, an annealing temperature of 60 °C and an extension time of 20 seconds. PCR products were purified by ethanol precipitation, by adding a final concentration of 300 mM NaAc (in-house, 3M stock) and 66% ethanol (Boom, 84028185) to the PCR reaction, incubating at −80 °C for 1 hour and centrifuging at 14,000 g for 30 minutes at 4 °C. The supernatant was removed and the DNA pellet was air-dried before resuspending in a total volume of 200 µL clean water. The purity of the DNA adapter was assessed on agarose gel (Invitrogen, 16500500) and the concentration was determined using a NanoDrop™ 2000 (Thermo Scientific).

The DNA probe was assembled in two steps. First, DBCO-PEG4-maleimide linker (Sigma, 760676; 2 nmol/µL in DMSO) was conjugated to the azide-labelled DNA adapter by combining 5 nmol linker (2.5 µL) with 1 nmol adapter (200 µL, ∼2,000 ng/µL) and incubating for 2 h on a rotor at room temperature. The adapter–linker was purified through an Amicon Ultra Centrifugal Filter, 3 kDa MWCO, washing at least three times (top up to 500 µL with 1X PBS, pH 7.4, 14,000 g, 10 min, room temperature) and recovered by inverting the filter into a fresh vial and spinning at 1,000 g for 2 min. Second, the adapter–linker was conjugated to the peptide (1 nmol/µL in 1X PBS, pH 7.4) via maleimide–cysteine reaction by combining 2 nmol peptide (2 µL) with the adapter–linker and incubating for 1.5 h on a rotor at room temperature, followed by overnight incubation at 4 °C. The coupled DNA probe was purified through an Amicon Ultra Centrifugal Filter, 10 kDa MWCO (Merck, UFC5010), using the same wash procedure. Clean DNA probes were aliquoted and stored at −20 °C. Probe quality and coupling efficiency were assessed as described for fluorescent probes with the following adaptation: To emphasize the mobility shift in the coupled vs. uncoupled DNA, the DNA fragment was shortened using restriction digestion with HpyCH4III enzyme (NEB, R0618S) in CutSmart Buffer for 1h at 37°. The digested DNA or DNA probe were then run on polyacrylamide gel as described above.

#### Image acquisition

##### Live-cell imaging

For live-cell imaging experiments related to figures 1-3, cells were seeded on a 96 Well Square Black Glass Bottom (Ibidi, 89627-90) at ∼40% confluency 1-2 days prior to imaging. Immediately before imaging, the culture medium was aspirated and exchanged for pre-warmed (37 °C) imaging medium (CO2-independent Leibovitz’s L-15 Medium (Gibco, 11415064) supplemented with 10% FBS and 1% Pen/Strep). All live-cell imaging was conducted at 37 °C. As a segmentation marker, cells were stained with either a plasma-membrane dye (CellMask Orange Plasma Membrane Stain, Invitrogen, C10045) or a whole-cell viability dye (Calcein AM Viability Dye, eBioscience, 65-0853-78) for 10 min before the start of image acquisition, at the concentrations recommended by the vendors.

##### Microscopes

Imaging was carried out on one of three systems: 1) a Nikon TI inverted microscope running NIS Element Software (Nikon), fitted with a perfect focus system, a Yokagawa CSU-X1 spinning disc and an iXon Ultra 897 EM-CCD camera (Andor) and a motorized piezo stage; 2) a Nikon TI2 inverted microscope running NIS Element Software, fitted with a perfect focus system, a Yokagawa CSU-X1 spinning disc and a Prime 95B sCMOS camera (Photometrics) and a standard motorized stage; or 3) a Nikon TI2 inverted microscope with a large field-of-view X-light V3 spinning disc (Crest Optics), an ORCA-Quest2 camera and a motorized piezo stage. Each of the three microscopes was fitted with a temperature-controlled box. Images were acquired using the following objectives: Plan Apo 100x Oil, NA 1.45 (Figures 1b, 1c, 1f, 4b), Plan Apo 40x Air, NA 0.95 (Figures 2b, 2j, 4c, 4f, Supplementary Figure 2b), Plan Apo 20x Air, NA 0.75 (Figures 2d, 3d, Supplementary Figure 1c).

##### Waldo barcode readout in imaging

Barcodes were read out iteratively across sequential staining rounds. In each round, cells were stained with the fluorescent probe for 1.5 minutes at 1:400 (from stock concentration) in imaging medium (see ‘Live-cell imaging’) and then imaged while in staining solution. The fluorescence was subsequently removed by adding a nuclease mix (1 µL Nuclease P1, 10 µL nuclease buffer and 90 µL L15; P1 Nuclease NEB M0660S) and incubating for 2 minutes, after which cells were washed once with 500 µL imaging medium. This cycle of staining, imaging, nuclease cleavage and washing was repeated for each successive fluorescent probe until all barcodes had been read out.

##### Photoactivation of selected cells/ROI

To photoactivate specific cells or areas of interest during live-cell imaging medium was removed and refreshed with imaging medium, supplemented with photoactivatable far-red HaloTag dye (PA-JF646) ^23^ at 0.5 µM final concentration and incubated for at least 5 min. To photoactivate an area of interest, the selected cells were imaged at 100% laser power (405 nm) for 10s at a single z position. The photoactivated PA-JF646 emits in the far-red channel. Photoactivation was followed by a round of image acquisition in the green, red and farred channels, to ensure that the correct cells were photoactivated and to determine the efficiency of photoactivation.

#### FACS sorting

To collect cells for FACS sorting and subsequent processing for single-cell sequencing, the medium was removed and cells were washed once with 200 µL 1X PBS, pH 7.4. Cells were then detached by adding 50 µL of enzyme-free Cell Dissociation Buffer (Gibco, 13151014) per well of a 96-well plate and incubating for 5 minutes. Cells were collected in 250 µL of ice-cold Leibovitz’s L-15 Medium with 10% FBS and 1% PenStrep (L15+/+) and centrifuged for 1.5 minutes at 800 g at 4 °C. These settings were used for all subsequent centrifugation steps, and cells and buffers were kept on ice from this point onwards. After centrifugation, the supernatant was removed and cells were resuspended in 250 µL ice-cold staining buffer (SB; L15+/+ with sequencing-probes, each at 1.25 ng/µL final). Cells were incubated with SB on ice for 15 minutes, then centrifuged and resuspended in 250 µL ice-cold L15+/+ as a wash. After another round of centrifugation, the supernatant was removed and cells were resuspended in 400 µL ice-cold L15+/+. Cells were pipetted through a Cell Strainer Snap Cap into a Falcon 5 mL Round Bottom Polypropylene Test Tube (Fisher Scientific, 10314791). Single cells were sorted on a BD Influx, BD FACSJazz or Beckman Coulter CytoFLEX SRT cell sorter into 384-well PCR plates (Bio-Rad, HSP3831) containing 50 nL CEL-Seq2 primer (1500 nM, as described in Rooijers et al.^26^, see ‘Sequences, primers and adapters’) and 5 µL mineral oil (Sigma, M8410) per well. Mineral oil was dispensed into plates pre-sorting using a Tecan Freedom EVO robot (Tecan), followed by dispensing of a unique CEL-Seq2 primer into each well using a Mosquito HTS robot (TTP Labtech). After sorting, plates were sealed with aluminium covers (Greiner, 676090), spun for 2 minutes at 2000 g at 4 °C and stored at −80 °C before further processing.

#### Waldo sample processing for single-cell sequencing

For Waldo sample processing, the steps to generate cDNA as well as the subsequent proteinase K digestion largely follow previously described protocols of Markodimitraki et al.^55^, including lysis, reverse transcription, second strand synthesis and proteinase K digestion. Below, we first summarize these shared steps and then describe the subsequent Waldo barcode readout steps specific to this study. Briefly, 384-well PCR plates with sorted cells were processed using a Nanodrop II robot at 12 psi pressure (BioNex) for adding all solutions. Indicated volumes are per well. Between handling and before robotic dispensions, plates were always spun for 2 minutes at 2000 g at 4 ⁰C. Plates were kept on coolblocks on ice in between robotic dispensions and incubations. After thawing plates (with sorted cells) on ice and plate centrifugation, 50 nL lysis mix (0.15% Igepal CA-630 (Sigma, I8896), 1:250,000 ERCC RNA spike-in (Ambion, Fischer Scientific, 13499968), 2 mM dNTPs (equimolar mix of dATP, dGTP, dTTP, dCTP, Invitrogen, 10297018) in RNAse-free water) was added to a total volume of 100 nL per well (including previously dispensed 50 nL CEL-Seq2 primer). Plates were incubated in a PCR machine for 5 minutes at 65 ⁰C, cooled to 4 ⁰C and transferred to ice. Immediately, 75 nL reverse transcriptase mix (4.67 U/μL RNaseOUT Recombinant Ribonuclease Inhibitor (Invitrogen, 10777019), 23.33 U/μL SuperScript II Reverse Transcriptase (Invitrogen, 18064022) in 2.33X manufacturer-supplied first strand buffer and 23.33 mM DTT in RNAse-free water) was added to a total volume of 175 nL. Plates were incubated in a PCR machine for 2 hours at 42 ⁰C, cooled to 4 ⁰C and transferred to ice. The PCR machine was then heated to 70 ⁰C and plates were incubated for 10 minutes at 70 ⁰C, cooled to 4 ⁰C and transferred to ice. 925 nL of second strand mix (270 μM dNTPs, 0.08 U/μL E.coli DNA ligase (NEB, M0205L), 0.27 U/μL DNA polymerase I (Invitrogen, 18010025), 0.016 U/μL RNase H (Invitrogen, 10378002) in 1.19X second strand buffer (Invitrogen, 10124942) in RNAse-free water) was added to a total volume of 1100 nL per well. Plates were incubated in a PCR machine for 2 hours at 16 ⁰C, cooled to 4 ⁰C and transferred to ice. 250 nL of proteinase K mix (5.2 mg/mL proteinase K (Roche, 3115879001) in 5.4X CutSmart buffer (NEB, B6004S in RNAse-free water)) is added to a total volume of 1350 nL per well. Plates were incubated in a PCR machine for 10 hours at 55 ⁰C, 20 minutes at 80 ⁰C, cooled to 4 ⁰C and transferred to ice. Next, 100 nL of variable spike-ins (either unconjugated probe DNA adapters, at 5,000-200,000 molecules per well, or peptide-coupled probe DNA adapters from peptides that were not used in the Waldo barcoding at 5,000-200,000 molecules per well) and 50 nL of constant spike-ins (either unconjugated probe DNA adapters at either 25,000, 50,000 or 100,000 molecules per well, or peptide-coupled probe DNA adapters from peptides that were not used in the Waldo barcoding at 25,000, 50,000 or 100,000 molecules per well) were added to a total volume of 1500 nL per well. 100 nL of EagI digestion mix (3 U/µL EagI-HF (NEB, R3505L), 2.5X CutSmart buffer in RNAse-free water) was added to a total volume of 1600 nL per well. Plates were incubated in a PCR machine for 3 hours at 37 ⁰C, cooled to 4 ⁰C and transferred to ice. 100 nL of cell-specific sample-adapter (110 nM, see section ‘Sequences, primers and adapters’) was added using a Mosquito HTS robot (TTP Labtech) to a concentration of ∼5.5 nM during ligation. 300 nL of ligation mix (1.08 U/µL T4 DNA ligase (Roche, 10799009001) with 6.66X of manufacturer-supplied Ligase Buffer in RNAse-free water) was added to a final volume of 2000 nL. Plates were incubated for 16 hours at 16 ⁰C, 10 minutes at 65 ⁰C, cooled to 4 ⁰C and transferred to ice. Safe-stopping points to store plates long-term during processing are after proteinase K digestion or after sample-adapter ligation.

#### Library preparation

Preparation and sequences of the uniquely-barcoded CEL-Seq2 primer sequences (set of 384 barcodes) are previously described in Rooijers et al.^26^. The preparation and sequences of the uniquely-barcoded double-stranded sample-adapters (top and bottom oligos, set of 384 barcodes) are previously described in Lochs et al.^24^, there referred to as SBC adapter sequences.

Primers used for library preparation are as follows, with sequences from 5’ to 3’ (ordered from IDT, with N as hand-mixed random bases): Random HexRT – GCCTTGGCACCCGAGAATTCCANNNNNN; EnrichRT – GCCTTGGCACCCGAGAATTCCACTAATTCACTCCCAACGAAG, the underlined sequence binds to receptor mRNA 3’UTR; RPi (example, each RPi has a unique index – see Markodimitraki et al.^55^) – CAAGCAGAAGACGGCATACGAGATCGTGATGTGACTGGAGTTCCTTGGCACCCGAGAA TTCC*A, in which * indicates a phosphorothioate bond that protects the DNA from exonuclease activity; RP1 - AATGATACGGCGACCACCGAGATCTACACGTTCAGAGTTCTACAGTCCG*A; BC-PCR-fw – TTCAGAGTTCTACAGTCCGACGATC; BC-PCR-rev – GTGACTGGAGTTCCTTGGCACCCGAGAATTCCACTCACCAGTCACAGAAAAGCATCTT;

##### Total IVT

After Waldo plate processing (as described in ‘Waldo sample processing for single-cell sequencing’, all cells with non-overlapping sample (single-cell) barcodes (generally a full 384-well plate) were pooled for in vitro transcription (IVT), generally following the library preparation protocol as previously described in Markodimitraki et al. and Lochs et al^.55,24.^ First, to pool cells, plates were spun upside down in a clean VBLOK200 Reservoir (ClickBio) for 4 minutes at 300 g, before collecting all of the water phase with residual mineral oil in one tube. Residual mineral oil was removed by spinning the sample for 2 minutes at 2000 g and transferring the liquid phase to a clean tube, which was repeated three times. After pooling, samples were incubated for 10-20 minutes with 0.8 volume CleanNGS magnetic beads (CleanNA, CPCR-0050), diluted 1:10 in bead binding buffer (20% PEG 8000, 2.5 M NaCl, 10 mM Tris–HCl, 1 mM EDTA, 0.05% Tween 20, pH 8.0 at 25 °C)). Samples were placed on a magnetic rack (DynaMag™-2, ThermoFisher, 12321D) to wash beads two times with 80% ethanol before allowing beads to dry before resuspending in 9 µL water. 1 µL of this material was transferred to a separate tube for subsequent Probe DNA PCR (see section ‘Probe DNA PCR’). IVT was performed by adding 12 µL IVT mix from the MEGAScript T7 kit (Invitrogen, AM1334) and incubating in a PCR machine for 14 hours at 37 ⁰C, cooling to 4 ⁰C and transfer to ice. Samples were first eluted from the beads and subsequently cleaned with 0.8 volume CleanNGS magnetic beads (undiluted) and three 80% ethanol washes, before eluting in 22 µL water and placing samples on ice. Next, the RNA was fragmented by adding 5.5 µL fragmentation buffer (200 mM Tris-acetate pH 8.1, 500 mM potassium acetate, 150 mM magnesium acetate) and incubating samples at 94 ⁰C for 90 seconds before immediately transferring to ice. The reaction was stopped by adding 2.75 µL STOP buffer (0.5 M EDTA pH 8). After transferring samples back to room temperature, samples were cleaned with 0.8 volume CleanNGS magnetic beads (undiluted) and three 80% ethanol washes, before eluting in 8.5 µL water – the clean amplified RNA (aRNA) sample. aRNA quality is evaluated on a Bioanalyzer RNA 6000 Pico Kit chip (Agilent, 5067-1513) and concentrations were measured using a NanoDrop™ 2000. 1 µL of this material was transferred to a separate tube for subsequent mRNA barcode enrichment (see section ‘mRNA barcode enrichment IVT’). Library preparation was subsequently performed as described previously53, using 5 µL of aRNA. 1 µL of 20 µM Random hexRT primer and 0.5 µL of 10 mM dNTP mix are added directly to the aRNA and the sample is incubated in a PCR machine for 5 minutes at 65 ⁰C, cooling to 4 ⁰C and transfer to ice. 2 µL of First Strand buffer, 1 µL DTT (0.1M), 0.5 µL SuperScript II reverse transcriptase and 0.5 µL RNAseOUT are added and the sample is incubated in a PCR machine for 10 minutes at 25 ⁰C, 1 hour at 42 ⁰C, cooling to 4 ⁰C and transfer to ice. Next, 2 µL of indexed RPi primer (10 μM) is added together with 25 µL NEBNext High Fidelity PCR Master Mix (NEB, M0541S), 2 µL RP1 primer (10 μM) and 10.5 µL DNase-free water. Library PCR is performed with 8 to 11 PCR cycles, depending on aRNA yield, according to the following protocol: 98 ⁰C for 30 seconds; 8-11x (98 ⁰C for 10 seconds, 60 ⁰C for 30 seconds, 72 ⁰C for 30 seconds); 72 ⁰C for 10 minutes. After PCR, samples are cleaned twice with 0.8 volume CleanNGS magnetic beads (undiluted) and finally eluted in 8-12 µL DNase-free water. Library quality is evaluated on a Bioanalyzer High Sensitivity DNA Analysis chip (Agilent, 5067-4626) and concentrations were measured using a Qubit dsDNA Quantification Assay (Invitrogen, Q32851) on a Qubit 3.0 Fluorometer (Invitrogen). Libraries were run on an Illumina NextSeq500 platform (high output 2x75 bp) or an Illumina NextSeq2000 platform (high output 2x100 bp), at an estimated output of 1.0-2.0x10^5^ read-pairs per cell.

##### mRNA barcode enrichment from IVT samples

1 µL of clean aRNA (see ‘Total IVT’ section) was diluted in RNAse-free water to 5 µL total, which was used for library preparation. The protocol is largely identical to the library preparation of the ‘total IVT’ (as described in the previous section), but the hexRT primer is replaced by a specifically-designed primer that binds to the receptor mRNA (EnrichRT primer), using 1 µL of 20 µM primer. The subsequent steps remain as described, with 11 PCR cycles and final elution of the DNA library in 8 µL to ensure a high concentration.

##### Probe DNA readout by PCR

1 µL of cleaned and pooled pre-IVT material (see ‘Total IVT’ section) was used for Probe DNA PCR. The PCR is performed using with Phusion High-Fidelity DNA Polymerase (NEB M0530L), with BC-PCR forward and reverse primers at 0.5 µM final concentration, 25 cycles and the following PCR incubations – 98 ⁰C for 30 seconds; 25x (98 ⁰C for 10 seconds, 58 ⁰C for 30 seconds, 72 ⁰C for 45 seconds); 72 ⁰C for 5 minutes. After PCR, samples were run on a 2% agarose gel to assess product quality and the correct band was purified using a QIAquick Gel extraction kit (QIAGEN, 28704), eluting in 30 µL DNase-free water. 1 µL of the purified sample was diluted to 21 µL total using DNase-free water. Next, 2 µL of indexed RPi primer (10 μM) was added together with 25 µL NEBNext High Fidelity PCR Master Mix and 2 µL RP1 primer (10 μM) for subsequent library PCR (as described in the ‘Total IVT’ section). The subsequent steps remain as described in section ‘Total IVT’, with 8 PCR cycles and final elution of the DNA library in 15 µL. Libraries were sequenced at an estimated output of 2.0-2.5 x 10^4^ read-pairs per cell.

#### Experiments with monoclonal cell lines expressing various Waldo Barcodes (Figures 2f-h, 3j-m)

To generate cell lines carrying defined Waldo barcodes, HeLa cells were transduced with a varying set of four different Waldo receptors, yielding ten polyclonal lines each expressing a distinct subset of Waldo receptors. Two weeks after transduction, single cells were sorted by FACS into individual wells to establish 15 monoclonal cell lines derived from the parental 10 polyclonal lines. Therefore, 10 out of the 15 monoclonal cell lines contain unique combinations of receptors while 5 monoclonal cell lines have overlapping receptor combinations with other clones but at different expression levels.

#### Proof-of-concept experiment (Figure 4 and Suppl. Fig. 4,5)

##### Generation of Waldo cell line

RPE-1 cells were engineered to stably express H2B-mScarlet and superfolderGFP-NLS (GFP-NLS). The constructs were introduced by lentiviral transduction followed by generation of a monoclonal cell line. The monoclonal cell line was then transduced with the lentiviral barcode library consisting of 10 Waldo receptors.

##### Experimental design

To induce chromosome missegregation and aneuploidy, cells were treated with 62.5 nM Cpd5 (Mps1 inhibitor; as described in^35^) for approximately 22 hours. DMSO-treated cells served as a negative control. Three independent biological replicates were performed. Per replicate, two wells of Cpd5-treated cells were imaged and subsequently processed for the full Waldo barcode readout workflow. One well with DMSO treated cells was imaged and analyzed for live-cell phenotyping but not further processed for single-cell sequencing and Waldo matching. Additional cells that were not imaged were sorted and processed for sequencing in each biological replicate: a well with Cpd5-treated cells and a well with DMSO-treated cells, both were used for sequencing quality assessment and validation.

##### Live-cell imaging

Cells were seeded in 96-well glass-bottom plates approximately 2 h before imaging, in medium containing either Cpd5 or DMSO. Time-lapse imaging was performed on a Nikon TI2 inverted microscope with a large field of view X-light V3 spinning disc (Crest Optics) and an ORCA-Quest2 camera and a motorized piezo stage (see “Microscopes” section) and were imaged using a 40x air objective (NA 0.95). Images were acquired every 12 minutes for 20–22 hours in two channels using 555 nm (H2B-mScarlet) and 488 nm (GFP-NLS) excitation lasers. Six z-planes were acquired at each timepoint with 1.5 µm spacing. Images were recorded in 16-bit with 2×2 pixel binning.

##### Sequential barcode staining and photoactivation

Following live-cell imaging, cells were subjected to 10 sequential rounds of Waldo probe staining and image acquisition as described in the ‘Waldo barcode readout in imaging’ section. Photoactivation was performed at the final timepoint as described in the “Photoactivation of selected cells/ROI” section. Cells were then collected, incubated with sequencing probes, and sorted as described in the “FACS sorting” section. Sorted cells were processed for library preparation as described in the “Waldo sample processing” and “Library preparation” sections.

##### Expected aneuploidy rate for dividing and non-dividing cells

To quantify the expected aneuploidy rate of dividing cells, we reasoned as follows. Aneuploidy in this experiment has two possible origins. First, a fraction of cells is called aneuploid at the start, independent of their phenotype during the experiment. We measured this background rate in DMSO control cells: ∼12% (Fig. 4e,g). Second, cells that divided in the presence of Cpd-5 can acquire aneuploidy through chromosome missegregation at that division.

A dividing cell is therefore aneuploid if either of these applies. Importantly, the two are not additive: a cell that was already aneuploid before division cannot become aneuploid again, so newly acquired aneuploidy can only occur in the ∼88% of cells that were called as near-diploid to begin with. If d is the fraction of initially near-diploid dividing cells that missegregated a chromosome, the expected aneuploidy rate of dividing cells is: 12% + 88% × d Non-dividing cells, by contrast, are simply aneuploid at the background rate of ∼12%.

Since ∼50% of cells divided during the experiment (Fig. 4d), the aneuploidy rate of the whole imaged population is the average of these two groups. Setting this equal to the measured total (∼25%): 0.5 × 12% + 0.5 × (12% + 88% × d) = 25% This simplifies to 12% + 44% × d = 25%, so 44% × d = 13%, giving d = 13/44 = 29.5%. Substituting this back into the expression for dividing cells: 12% + 88% × 29.5% = 12% + 26% = 38%. The expected aneuploidy rate is therefore ∼38% for dividing cells and ∼12% for non-dividing cells.

### Computational methods

#### Calculation of barcode diversity

To estimate the theoretical barcode diversity using combinatorial Waldo receptor expression and to understand how different features of receptor expression impact barcode diversity, we performed simulations across a broad parameter space of cell population size, numbers of barcode receptors, receptor expression distribution and measurement noise. For each condition, a synthetic expression matrix of N cells × d receptors was generated, drawing each receptor’s expression independently from one of nine parameterized receptor expression distributions. These included two continuous unimodal distributions (uniform U[0,1]; normal μ = 0, σ = 2), three bimodal distributions (mixtures of two Beta distributions, representing receptors with two preferred expression states), and atwo binary Bernoulli distributions (p = 0.3 or 0.5) as the limiting ON/OFF case. To capture a receptor that is absent in some cells but expressed at varying levels in others, we used a spike-and-slab distribution: a point mass at zero (probability p_spike = 0.2) mixed with a Beta-distributed slab (Beta(3,3) or Beta(5,3)), with a small uniform jitter (0.01) to avoid exact zeros. Each receptor column was then min-max normalized to [0,1]. Simulations spanned all combinations of N ∈ {50, 100, 200, 400, 500, 1000, 2000, 5000, 10 000, 20 000, 50 000} and d ∈ {6, 8, 10, 12, 14, 16, 18, 20, 25, 40} (110 combinations per distribution), each simulation was replicated three times with different random seeds.

To assess whether two barcodes that were similar could still be distinguished, a key parameter that sets overall barcode diversity, each cell i was treated as a multivariate Gaussian over d receptor dimensions, with mean equal to its expression vector μᵢ ∈ [0,1]ᵈ and isotropic covariance Σ = σ²I, where σ is the measurement noise. Confusion between cells i and j was measured by the Bhattacharyya overlap coefficient, which for equal-covariance Gaussians reduces to

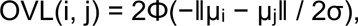

where Φ is the standard normal CDF and ǁ·ǁ the Euclidean distance. OVL ranges from 0 (fully distinguishable) to 1 (identical) and reflects the probability that noisy measurements of two cells cannot be reliably told apart. For each cell, OVL was computed against all N − 1 others and the cell classified, given a threshold τ, as unique (OVL < τ for all j), non-unique (OVL ≥ τ for all j) or ambiguous (OVL < τ for some but not all j). The three fractions sum to 1; the primary outcome was the fraction of uniquely barcoded cells. Noise and threshold values were swept across σ ∈ {0.01, 0.05, 0.1, 0.2, 0.5} and τ ∈ {0.001, 0.01, 0.05, 0.1}; τ = 0.05 was used as the default for all main-text results unless stated otherwise. For large populations (up to N = 50 000), the O(N²) pairwise distances were computed in row chunks using BLAS-accelerated matrix multiplication (tcrossprod()) to avoid materializing the full distance matrix, capping peak memory at ∼2–3 GB per task. Each condition ran as an independent SLURM array job. Results from the three replicates were pooled per parameter combination (distribution, N, d, σ, τ) to compute the mean, standard deviation and standard error of each fraction. All simulations and analyses were performed in R (≥ 4.2) using tidyverse, furrr, future, data.table and ggplot2.

#### Image analysis of Waldo datasets

##### Quantification of Waldo receptor and probe staining intensities (Figures 1D,1E, 2D-H)

If multiple z-slices were acquired, maximum intensity projection was performed for further quantification in 2D. Cells were segmented based on either whole-cell staining or membrane staining (see ‘Image acquisition: Live cell imaging’) using cellpose 2.0 with cyto model and mean intensities were extracted for each segmented cell.

##### Live-cell imaging datasets

Live-cell imaging field of views (FOVs) were corrected for inhomogeneity in illumination and adjacent FOVs were stitched together using NIS elements software. Using these large, intensity-corrected, stitched FOVs cells were then manually annotated for phenotypes such as cell division, nuclear deformation, micronuclei and micronuclear rupture. Annotations were saved to the coordinates of the corresponding cells in the last timepoint of the live-cell imaging datasets to allow linking of cellular phenotypes to their (live-cell) Waldo barcode, and eventually to the single-cell sequencing data for the respective cell.

##### Fluorescent barcode readout

For each imaging round of the fluorescent barcode readout, a separate nd2 imaging file was created. After finishing the complete barcode readout, images from barcode rounds were concatenated into a single nd2 file, corrected for illumination artefacts, maximum-intensity projected and stitched per timepoint using NIS elements software to yield one barcode readout timelapse containing the stitched images per well. Using this barcode timelapse data, cells were then segmented on the whole-cell marker (see “Live-cell imaging”) using Cellpose^57^ (cyto-SAM model). Cells were then tracked over timepoints using trackastra^58^. The tracked output segmentations were then manually curated. Single-cell intensities for the Cy5 (barcode) channel were then extracted and the median intensity (median pixel value per cell segmentation) was used for all downstream steps. To correct barcode intensities for local background, cell segmentations were locally expanded by up to 50 pixels into background regions containing no other segmentations. The 5th-percentile from the newly expanded region was then used to calculate the local background and was subtracted from the median single-cell receptor expression intensity extracted earlier. Negative values were set to zero, resulting in a distribution resembling a spike-slab distribution. The non-zero values (slab) were then log2-transformed and mad-scaled.

#### Processing of sequencing data

Waldo, scKaryo and scKaryo&T libraries were processed with a common workflow, based on the workflow described in Markodimitraki et al.^55^, followed by experiment-specific read alignment and quantification. For the experiments in figures Suppl. Figure 4 reads were aligned against the standard human reference genome hg19 (GRCh37), whereas for the proof-of-concept experiments reads were aligned against a cell-line-matched, haplotype-resolved RPE-1 reference to enable better copy-number calling; this difference propagates into distinct pipeline steps in the alignment and quantification. Shared steps are described first, then the procedures specific to the validation experiments (Figures 3E–M, Suppl. Figure 4) and proof-of-concept experiments (Figure 4, Suppl. Figures 5–6).

##### Common processing

###### Demultiplexing

Reads were demultiplexed on the barcode at the start of R1 using a reference barcode sequence list, allowing no mismatches. For Waldo (CEL-Seq2 reads; karyo-reads; Waldo barcodes), scKaryo&T (karyo-reads; CEL-Seq2) and scKaryo (karyo-reads) libraries, the reference contained either single adapter barcodes or a concatenation of the library’s barcodes. The UMI, also at the start of R1, was appended to the read name. Each library was processed across up to three modalities: CEL-Seq2 mRNA reads, scKaryoSeq reads, and Waldo barcode reads.

###### Detection of Waldo probe DNA barcode reads

Demultiplexed reads were subject to probe DNA barcode detection. First, per single-cell sample, reads were scanned for the constant linker sequence (GGCCGCCGTTCTAC) that has a fixed spacing from the probe DNA barcode and UMI, allowing 1 mismatch. Reads containing the linker sequence were kept for probe DNA barcode detection and their UMI sequence was stored for later read deduplication. The probe DNA barcode was detected at a constant distance from the linker sequences, allowing 1 mismatch. For each sample, the abundance of each probe DNA barcode was finally determined as the number of total probe DNA barcodes that were detected across demultiplexed reads.

###### Detection of Waldo mRNA barcode reads

CEL-Seq2 reads were aligned using hisat2 (v. 2.1.0)^59^ with the following parameters: “--mp ‘2,0’ --sp ‘4,0’”, using a newly created reference genome consisting of the receptor-construct sequence inserted into the human reference genome. All reads that were successfully aligned to the receptor-construct reference were scanned for the linker sequence just before the receptor-specific barcode insertion site, without allowing for any mismatches. For reads where the linker sequence was identified, the receptor-specific barcode was extracted and added to a count table, together with the UMI of the read. This table was later used to calculate unique receptor mRNA read counts for single cells.

###### Cell quality filtering

For each single-cell sequencing modality (scRNA-seq, scKaryoSeq), cells were removed if they contained low read counts, based on the bimodal distribution of unique aligned read counts. The threshold was determined as the local minimum between the two Gaussian distributions using a Gaussian-mixture model, then visually inspected and adjusted if necessary. Waldo mRNA barcode and CEL-Seq2 readouts were filtered on CEL-Seq2 counts; Waldo probe DNA barcode and scKaryoSeq readouts on karyo-read counts. Integrated analyses required cells to pass both scRNAseq and scKaryoSeq modality-specific filters.

##### Waldo barcode readout for validation experiments (Figures 3E–M, Suppl. Figure 6)

###### CEL-Seq2 data processing

Reads were aligned to hg19 (GRCh37) with hisat2 (v. 2.1.0)^59^ using “--mp ‘2,0’ --sp ‘4,0’”; only R2 was used, as R1 carried the sample barcode, UMI and poly-A tail. Alignments were converted to unique transcript counts per gene with custom scripts analogous to HTSeq’s^60^ htseq-count in “intersection_strict” mode.

###### scKaryoSeq data processing

A ‘C’ nucleotide, digested from karyo fragments by EagI during sample processing, was prepended to the reads, which were then aligned to hg19 (GRCh37) with bowtie2 (v. 2.4.1)^61^ using “--seed 42 --very-sensitive -N 1”. Alignments were converted to UMI-unique ‘CGGCCG’ counts by matching to known CGGCCG positions; reads not mapping to a known position or with mapping quality below 10 were removed. Depending on the cell line’s reported karyotype, two or three unique UMIs were allowed per CGGCCG position, and counts were binned at the desired resolution.

###### OE-value calculations

Observed-over-expected (OE) values were calculated as described previously^26^. For each bin i in cell j, OEᵢ,ⱼ = (nᵢ,ⱼ / mᵢ) × (Σmᵢ / Σnᵢ,ⱼ), where nᵢ,ⱼ is the observed uniquely mapped reads in segment i, Σnᵢ,ⱼ the total genome-wide in cell j, mᵢ the maximum expected unique reads in segment i, and Σmᵢ the maximum expected genome-wide.

##### Waldo proof-of-concept experiments (Figure 4, Suppl. Figures 4–5)

###### Reference genome and gene annotation

Data were aligned to the diploid RPE-1 reference genome (GIUNTAlab RPE1V1.1;^62^), with both haplotypes represented as separate contigs (chr1_hap1, chr1_hap2, …). Gene annotations were transferred from the hg38 Ensembl GTF to each haplotype independently using Liftoff^63^ (coverage and identity thresholds 0.95). The per-haplotype GTFs were merged and contig names remapped with custom scripts into a unified diploid GTF.

###### CEL-Seq2 data processing

Reads were aligned to the diploid RPE-1 genome with STAR^64^ (v. 2.7.10) using forward-strand specificity and minimum mapping quality 10 and assigned to genes with featureCounts^65^ (v. 2.1.1). UMI deduplication and count matrix construction used a custom script (build_count_matrix_v2.py), collapsing reads mapping uniquely to either haplotype or equally to both haplotypes of the same gene into one per-gene count per cell.

###### scKaryoSeq processing

Reads were aligned in paired-end mode with Bowtie2^61^ (v. ≥ 2.5.0) using --very-sensitive-local and -k 10; local alignment accommodated the ∼6 bp EagI adapter prefix on R1. Read pairs were classified as uniquely haplotype-1, uniquely haplotype-2, ambiguous-valid (equal best score to both haplotypes of the same chromosome), or discarded (classify_reads.py). UMI deduplication used a custom greedy algorithm with ±5 bp position tolerance. Retained reads were pooled per chromosome into ∼500 variable-width bins; bins with high cross-cell variability or mappability artefacts were blacklisted from quality metrics across QC-passing cells.

###### Copy number calling

Per-cell bin counts were normalised to counts per 1,000 reads and divided by the per-plate per-bin median across quality-passing cells. A chromosome was called aneuploid only when the within-cell interquartile range [Q25, Q75] of normalised values lay entirely above 2.25 (gain) or below 1.75 (loss). Chromosomes X (sex), 10 (constitutive CNV in RPE-1), 18 and 21 (insufficient non-blacklisted bins) were excluded. The aneuploidy score per cell was calculated as Σ|CN − 2| across included chromosomes.

###### scRNAseq integration, UMAP and cell cycle scoring

Gene expression was normalised to counts per million on QC-passing cells and log1p-transformed. Batch effects across experiments and plates were corrected with Harmony^66^. Cell cycle scores (S, G2/M) were computed with Seurat v5^67^. UMAP coordinates were computed on Harmony-corrected principal components.

###### Pathway activity analysis

Pathway activity was scored as the mean z-scored CPM across gene-set members (Suppl. Table 3).

#### Imaging–sequencing cell matching

Within an imaging well, the number of imaged cells substantially exceeded the number of cells that were successfully sorted into 384-well plates and survived sequencing quality control. Each sequenced cell must therefore be matched to its imaging counterpart within a much larger pool of candidates, most of which are not the true match. A direct quantitative comparison of barcode values between modalities is complicated by the fact that imaging and sequencing measure receptor levels on different scales. The matching algorithm therefore proceeded in two stages: first, the large imaging population was narrowed to a small candidate set based on which Waldo receptors were expressed—a binary pattern that is robust across modalities—and only within that reduced set were quantitative barcode scores compared to identify the single best match.

##### Composite sequencing barcode score

For each sequencing plate, barcode positivity was determined per cell and Waldo receptor using mRNA signal only. A two-component Gaussian mixture model was fit to the log1p-transformed UMI counts of each Waldo receptor, and the valley between the two components defined the threshold separating positive from negative calls. Probe-DNA signal was not used for this classification: a receptor not called positive by the mRNA threshold was treated as absent regardless of its probe-DNA level. For each cell and receptor called positive, a composite barcode score was computed by combining all available sequencing readouts. Each readout—up to four, comprising mRNA and/or probe-DNA reads from the IVT and/or enrichment libraries—was z-scored per source, receptor, and plate across the QC-passing cell population, and the resulting z-scores were averaged into a single composite score. This design accommodates experiments with different library combinations while integrating information from multiple readouts where available. The composite barcode score served as input for downstream matching to imaging barcodes.

##### Two-stage matching algorithm

Sequencing cells were matched to imaging cells separately within each well.

*Stage 1—candidate selection by soft Jaccard similarity.* For each of the 10 Waldo receptors, composite barcode scores (sequencing) and barcode intensity values (imaging) were classified as positive or negative using a threshold of zero. Among positive receptors, values were rank-transformed to (0, 1] separately within the sequencing and imaging populations; negative receptors were set to zero. For every sequencing–imaging cell pair, a soft Jaccard similarity was computed as

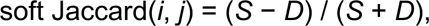

where *S* is the summed rank-normalised value across all 10 Waldo receptors for the pair, and *D* is their L1 distance in rank space. This generalises binary Jaccard similarity to continuous-valued data: shared absences contribute to neither *S* nor *D*, while mismatches involving high-ranking positive values are penalised more heavily than those involving low-ranking ones. For each sequencing cell, all imaging cells with a soft Jaccard score ≥ 0.4 were retained as Stage 2 candidates; if fewer than 5 imaging cells passed this threshold, the top 20 were kept instead, ensuring a non-empty candidate pool.

*Stage 2—matching by Pearson correlation.* Within each candidate pool, composite barcode scores and imaging barcode intensities were each transformed to approximate normality using a rank-based quantile transform applied separately per modality. Receptors called negative were assigned a fixed z-score (qnorm(0.01)), so that shared absences contributed consistently to the correlation regardless of differences in the negative rate between modalities. For each candidate pair, Pearson correlation was computed across the 10 Waldo receptors, and the imaging cell with the highest correlation was assigned as the match. The gap between the best and second-best correlation and the candidate pool size were retained as match-confidence diagnostics. In all downstream analyses, a post-hoc quality filter requiring at least three Waldo receptors with a positive barcode call was applied, removing matches with low barcode expression without altering the assignment itself.

